# Transient noradrenergic signals act as online controllers of perceptual computation

**DOI:** 10.64898/2026.10.21.760810

**Authors:** Supriya Ghosh, John H. R. Maunsell

## Abstract

Neuromodulatory signals influence perception, arousal and learning, but their effects on behavioral performance can arise from fundamentally different computations. A transient signal could alter a persistent operating state, control an ongoing computation, or provide a teaching signal that modifies future behavior. These mechanisms make distinct predictions about the time course of neuromodulatory influence on behavior within and across trials. Here we applied temporally randomized white-noise optogenetic activation of locus coeruleus (LC) noradrenergic neurons while monkeys performed a visual match-to-sample discrimination task. Computational simulations established that reverse correlation could distinguish the temporal signatures of the modeled persistent-state, controller and outcome-dependent teaching mechanisms based on behavioral choices. LC activation selectively increased perceptual sensitivity for contralateral stimuli during a narrow time window preceding sensory input, without a detectable change in decision criterion. Cross-trial effects were smaller than predicted by the persistent-state and teaching models, despite sufficient sensitivity to detect the predicted effects. After accounting for the measured LC spiking response to optical stimulation, we estimated that behaviorally effective LC activity occurred approximately 50 ms before sensory input. The temporal weighting of optical stimulation predicted contralateral perceptual outcomes on held-out individual trials. Together, these findings support a temporally and spatially selective controller-like contribution of transient LC activation to ongoing perception among the mechanisms tested. More broadly, they demonstrate how the temporal structure of causal effects can distinguish computational roles of neuromodulatory signals that are difficult to resolve from average behavioral performance.

## Introduction

The computational role of transient locus coeruleus (LC) noradrenergic signaling in perceptual behavior remains unresolved. Although LC noradrenergic neurons have been implicated in diverse cognitive functions, including perceptual sensitivity, attention, arousal, learning and behavioral flexibility ^1-7^, these observations leave unresolved a fundamental question regarding the computational role of a transient LC-noradrenergic event. A transient LC-noradrenergic event could alter a persistent operating state that influences multiple ongoing computations, transiently regulate the computation being performed at that moment, or provide a teaching signal that modifies future behavior through learning. In the persistent-state model, LC input directly changes an operating state independently of behavioral outcome. In the teaching model, LC input interacts with outcome feedback to update a policy-weight that influences subsequent behavior. Because these mechanisms can produce similar behavioral effects when averaged over time or across trials, distinguishing among them experimentally has remained difficult. Importantly, however, they make fundamentally different predictions about when the consequences of transient LC-noradrenergic activity are manifest and how long they last.

LC function has commonly been examined using sustained manipulations, perturbations aligned to predefined task events or trial-averaged measurements ^6-8^. Such approaches can establish whether LC influences perception or behavior, but provide limited information about when transient LC activity influences underlying computations and whether its influence persists after LC activity ends. Temporally precise perturbations provide an approach to identifying the computational policies implemented by transient LC-noradrenergic signaling, rather than simply establishing whether LC influences perception. In particular, resolving its causal influence both within the ongoing trial and across subsequent trials could distinguish mechanisms that appear similar when averaged over time.

Here we formalized three competing computational policies that assign distinct roles to transient LC-noradrenergic signaling (**Figs. 1a-c**). The models differ principally in what follows a brief LC event: a persistent change in operating state, a transient influence on the computation occurring at that moment, or an outcome-dependent update expressed on future trials. First, in a persistent-state model, brief LC activation displaces a latent state that sets the operating conditions for multiple ongoing computations, including sensory processing, memory retention and decision formation (**Fig. 1a**). Because the state evolves more slowly than the initiating LC event, its displacement can continue to influence sensory, memory and decision processes after LC activation has ended and can persist across multiple trials. This model is motivated by theoretical accounts linking LC-noradrenergic signaling to adaptive gain, uncertainty and internal-state regulation ^9,10^, together with evidence that LC activation can regulate sustained behavioral and distributed network states and that brief LC stimulation can produce cortical consequences that outlast the stimulus ^6,11,12^. In this framework, the enduring variable is the operating state, rather than the LC event itself.

**Figure 1.**
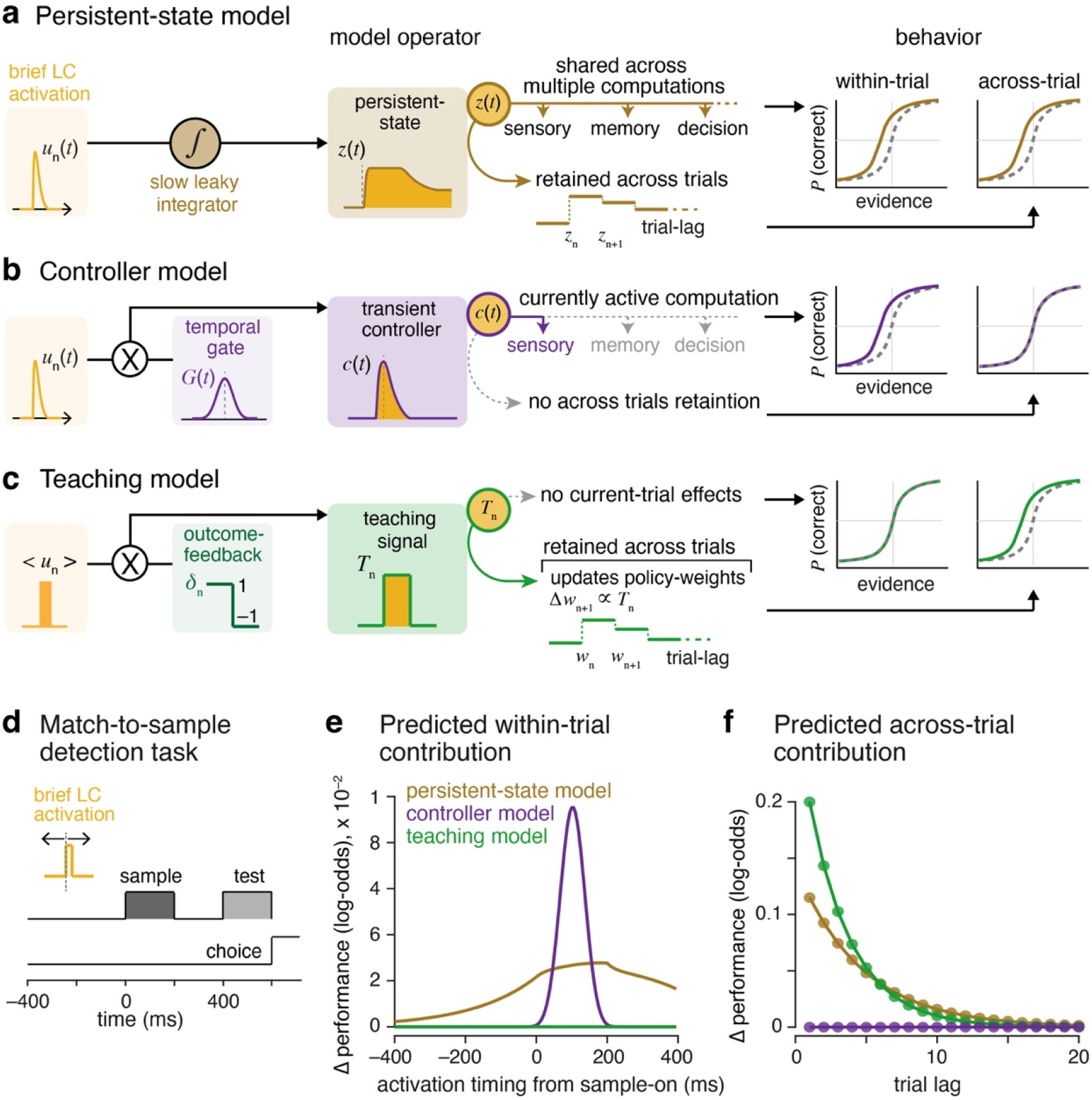
Competing computational hypotheses for how transient locus coeruleus activity influences perceptual performance. (**a**) Persistent-state model. Brief LC activation displaces a latent operating state, *z*(*t*), that sets the conditions under which multiple computations are performed, including sensory processing, memory retention and decision formation. Because this state evolves more slowly than the initiating LC event, its displacement can influence the remainder of the current trial and can be carried across trial boundaries, progressively decaying over subsequent trials. (**b**) Controller model. Brief LC activation influences behavior only when it coincides with a computation that is susceptible to LC modulation. This temporal susceptibility is represented by a computation-specific gate, *G*(*t*); LC input passing through the gate generates a transient controller signal, *c*(*t*), that modulates the computation active at that time. The controller signal is not retained across trials, so its effect is restricted to the ongoing trial. (**c**) Teaching model. Brief LC activation does not directly alter the choice being generated on the current trial. Instead, LC activation is combined with signed outcome feedback, *δ*_n_ (+1, correct; –1, error) after the choice, to generate a teaching signal, *T*_n_. The teaching signal has no effect on current-trial performance but updates a retained policy weight, representing a learned bias on subsequent behavior, whose influence progressively decays across future trials. (**d**) Delayed match-to-sample task and timing of brief LC activation. LC activation was considered at different times relative to sample onset, spanning pre-sample, sample, delay and test periods. (**e**) Predicted within-trial contribution to performance as a function of LC activation timing. The persistent-state model predicts a broad temporal contribution reflecting the dynamics of the operating state, whereas the controller model predicts temporally localized contributions determined by the position of the a priori temporal gate (shown for gates centered at +100 ms relative to sample onset). The teaching model predicts no within-trial contribution. (**f**) Predicted across-trial contribution to performance as a function of future-trial lag. The persistent-state model predicts a decaying contribution arising from retention of the operating-state displacement, whereas the teaching model predicts a decaying contribution arising from retention of the updated policy weight. The controller model predicts no across-trial contribution because the controller signal is not retained across trials. Contributions in **e** and **f** are expressed as changes in log-odds of correct performance relative to the corresponding no-activation condition.

Second, in a controller model, brief LC activation acts as a transient control input that regulates an ongoing computation without driving a persistent internal state (**Fig. 1b**). Its effect is expressed only during a computation-specific temporal window, represented by a gate *G*(*t*), during which the relevant process is susceptible to LC modulation. LC input passing through this gate generates a transient controller signal that acts on the computation active at that time, for example sensory, memory or decision processing. This hypothesis draws on the computational principle of online control, in which control signals regulate ongoing computations according to their state and timing ^13,14^, and on proposals that phasic LC activity rapidly reconfigures active target networks ^15^. Empirically, phasic LC activation can transiently enhance sensory cortical responses in a timing-dependent manner ^8,16^ and selectively improve perceptual sensitivity ^7^. In this framework, LC does not establish a lasting state; its behavioral consequence depends on whether the LC event coincides with a computation that is currently susceptible to modulation.

Third, in a teaching model, brief LC activation does not alter the perceptual computation generating the current choice. Instead, after the outcome of that trial becomes available, LC activation interacts with signed outcome feedback to produce an update that influences subsequent behavior (**Fig. 1c**). We represent this as a teaching signal that updates a retained policy weight, that is, a learned bias affecting future choices. This formulation is motivated by evidence that LC-noradrenergic activity carries signals related to reinforcement and prediction errors and can facilitate experience-dependent plasticity and subsequent learning ^5,17-19^. We formalized this possibility as an outcome-dependent update in which the influence of LC activation on future behavior is conditioned on signed trial outcome. Thus, unlike the first two models, the defining consequence of the teaching mechanism emerges only after feedback and is expressed on subsequent trials, without altering the perceptual computation generating the current choice. In the present well-learned task, this model tests whether transient LC activation produces outcome-dependent changes in subsequent performance under stable task contingencies; it does not test LC-dependent learning during acquisition or contingency changes.

The three hypotheses are motivated by broader theories of internal-state regulation, online control and learning, respectively, and we formulate them here as explicit computational operators with distinguishable temporal consequences. We reasoned that these differences could be resolved by treating transient LC activation as a system-identification signal. We therefore combined weak, temporally randomized white-noise (WN) optogenetic activation of LC with a visual match-to-sample task in monkeys. We used reverse-correlation analysis, extending approaches previously used to recover behavioral weighting functions from optogenetic WN perturbations ^20-22^, to estimate how LC activation contributes to perceptual performance as a function of its timing within the current trial and its history across preceding trials. We first tested whether the three computational policies generate identifiable signatures in simulated behavior and then applied the same framework to experimental choices. This approach allowed us to ask not simply whether LC influences perceptual behavior, but which computational policy best describes the causal influence of a transient LC-noradrenergic event.

## RESULTS

### Competing LC computational policies make distinct temporal predictions

We first asked how the three computational policies translate into experimentally distinguishable effects on perceptual behavior. We considered brief LC activation at different times during a delayed match-to-sample task, spanning sample encoding, memory delay and test processing (**Fig. 1d**). For each model, we quantified the predicted contribution of LC activation to perceptual performance, expressed as the change in log-odds of a correct response, both within the current trial and across subsequent trials. These two temporal dimensions provide the key diagnostic: persistence and control can both influence the current trial, whereas persistence and teaching can both influence future trials.

The three models produced distinct within-trial predictions (**Fig. 1e**). In the persistent-state model, brief LC activation displaced a latent state whose dynamics outlasted the activation itself and influenced multiple components of task processing. Consequently, the predicted contribution to current-trial performance was temporally broad. In the controller model, LC activation affected behavior only when it coincided with the temporal gate of the active computation. Its contribution was therefore temporally localized, and shifting the gate from sample onset to 100 ms after sample onset shifted the predicted contribution accordingly. The teaching model produced no within-trial contribution because its update is generated only after outcome feedback and therefore cannot affect the choice that produced that outcome.

The models also differed in how the consequences of LC activation extended across trials (**Fig. 1f**). In the persistent-state model, the residual displacement of the operating state was carried into subsequent trials with a progressively declining contribution to future performance. The controller state was reset at the trial boundary and therefore produced no across-trial contribution. In the teaching model, LC activation combined with outcome feedback to update a policy weight that influenced subsequent choices and decayed over future trials.

The models can therefore be distinguished by considering within- and across-trial effects jointly: persistent-state and controller mechanisms can both influence the current trial, but only the persistent-state mechanism produces effects that persist across trials; persistent-state and teaching mechanisms can both influence subsequent trials, but only teaching depends on trial outcome.

### Behavioral reverse correlation distinguishes competing LC computational policies

The theoretical predictions in **Figure 1** define the expected temporal consequences of LC activation under each computational policy, but behavior provides only stochastic choices from which these contributions must be inferred. We therefore tested whether these signatures could be recovered from simulated behavior using reverse correlation. Behavior was generated under each model using binary WN LC input and analyzed with the same generalized linear model (GLM) framework used for the experimental data (**Fig. 2**). The models produced similar overall performance (mean ±1 SEM, persistent-state, 68.03 ± 0.02%; controller, 66.04 ± 0.02%; teaching, 66.89 ± 0.03% correct; across 500 simulations, 8000 trials), allowing their temporal signatures to be evaluated at broadly comparable performance levels. Importantly, the temporal weights were not imposed on simulated choices: behavior was generated from each computational policy, and within- and across-trial weights were estimated from the resulting choices.

**Figure 2.**
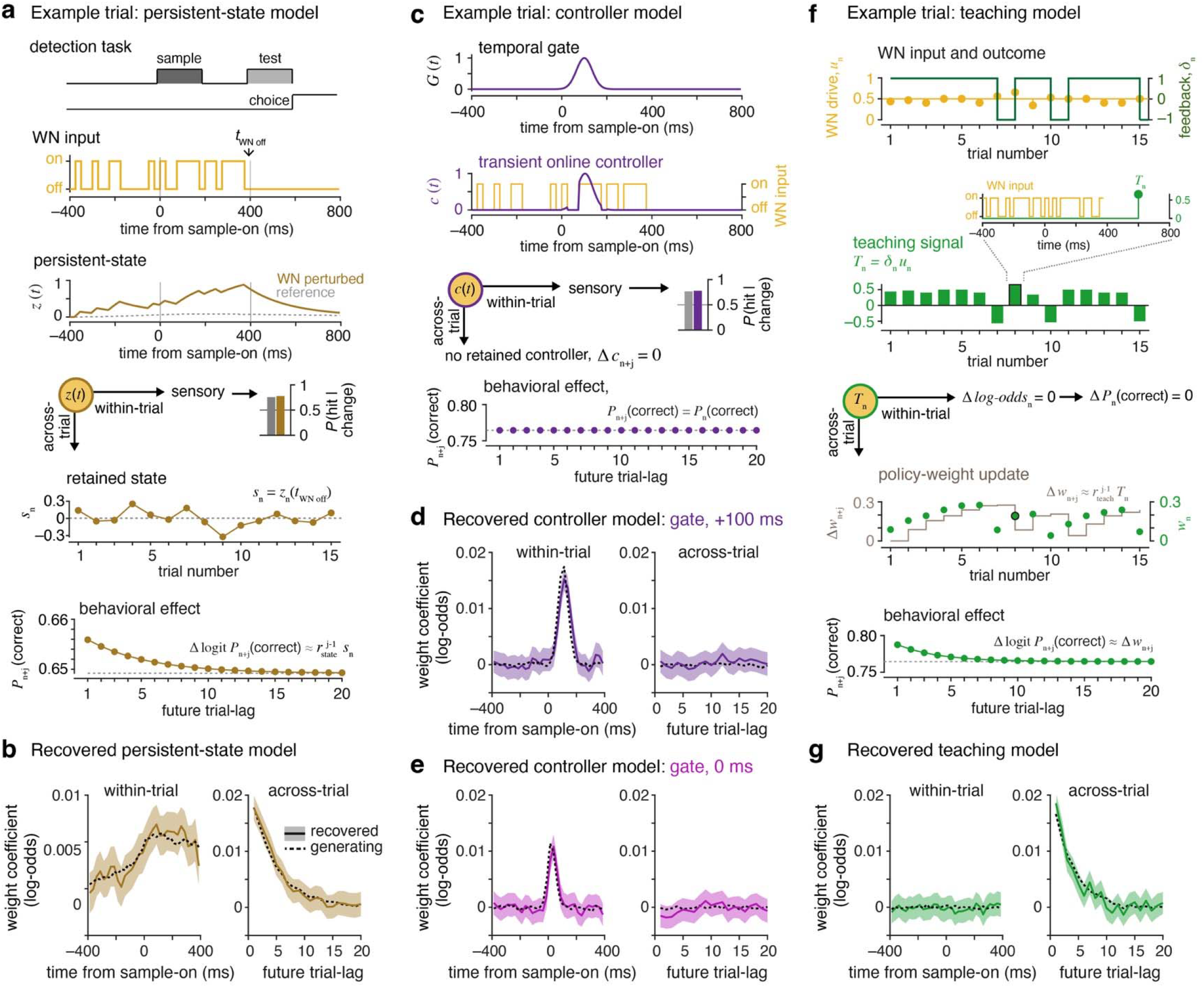
Behavioral simulations recover distinct within- and across-trial signatures of the three computational hypotheses. (**a**) Example simulation for the persistent-state model. Binary white-noise (WN) LC perturbations drive a latent operating state, *z*(*t*), that persists beyond individual perturbation epochs and modulates current-trial sensory processing, memory retention and decision-related processes, thereby altering the probability of a correct response. The WN-driven state remaining at perturbation offset, *s*_n_ = *z*_n_(*t*_WN off_), is retained across trial boundaries and produces a baseline-state displacement that decays over subsequent trials, resulting in a corresponding decaying influence on future-trial *P*(correct). (**b**) Within-trial (*left*) and across-trial (*right*) kernels recovered by GLM analysis of choices generated by the persistent-state simulation. Recovered kernels (solid) are shown with their uncertainty (shading, 95% confidence intervals) together with the theoretical generating model (dotted). The simulation recovers both the broad within-trial influence and the decaying across-trial influence predicted by the persistent-state model. (**c**) Example simulation for the controller model. LC perturbations are filtered by a temporally selective computational gate, *G*(*t*), producing a transient controller signal, *c*(*t*), that modulates sensory processing only during the currently engaged computation. The controller signal is not retained across trials, predicting a temporally localized within-trial effect but no across-trial effect. (**d-e**) GLM-recovered kernels for controller simulations with the temporal gate centered at +100 ms (d) or 0 ms (e) relative to sample onset. Shifting the gate correspondingly shifts the recovered within-trial kernel, whereas the across-trial kernel remains centered near zero. (**f**) Example simulation for the teaching model. Trial-level LC drive, *u*_n_, is combined with signed outcome feedback, *δ*_n_, to generate the teaching signal *T*_n_ = *δ*_n_ *u*_n_. The teaching signal has no effect on behavior within the trial on which it occurs [Δlogit *P*_n_ (correct) = 0], but updates a learned policy weight that influences subsequent trials. The teaching-induced policy-weight displacement, Δ*w*_n+j_, decays across future-trial lags and produces a corresponding decaying change in future *P*(correct). (**g**) GLM-recovered teaching model kernels. As predicted, the within-trial kernel remained near zero, whereas the across-trial kernel recovered the decaying influence of preceding LC-feedback events. Together, the simulations recovered the characteristic within- and across-trial signatures of the three computational mechanisms.

In the persistent-state model, WN LC input drove the latent operating state *z*(*t*), which modulated sensory processing, memory retention and decision-related processes within the trial (**Fig. 2a**). The LC-driven state remaining at perturbation offset was retained across trial boundaries, producing a baseline-state displacement that progressively decayed across subsequent trials. Reverse correlation recovered both predicted features: a broad within-trial contribution and an across-trial contribution that declined with trial lag, each closely following the theoretical prediction (**Fig. 2b**).

In the controller model, WN LC input was filtered by the temporal gate *G*(*t*), producing a transient controller signal that modulated the eligible within-trial computation without being retained across trials (**Fig. 2c**). With the gate centered 100 ms after sample onset, the recovered within-trial weights were localized to this period, whereas across-trial weights remained near zero (**Fig. 2d**). Shifting the gate to sample onset correspondingly shifted the recovered within-trial weights without introducing across-trial effects (**Fig. 2e**), demonstrating recovery of both the timing and trial specificity of controller operation.

In the teaching model, trial-level LC drive, *u*_n_, was combined with signed outcome feedback, *δ*_n_ (+1 correct, –1 error), to generate the teaching signal, *T*_n_ (**Fig. 2f**). The teaching signal did not influence behavior on the trial in which it occurred, but updated a policy weight whose influence on behavior decayed across subsequent trials. Accordingly, recovered within-trial weights remained near zero, whereas across-trial weights captured the predicted decaying influence of preceding LC-feedback events (**Fig. 2g**).

Together, the three models produced different within- and across-trial signatures under the same temporally varying LC input. The persistent-state model produces both within- and across-trial effects. The controller model produces a temporally localized within-trial effect without across-trial carryover. The teaching model produces outcome-dependent effects on subsequent trials. Neither within-nor across-trial effects alone fully distinguish the three mechanisms, but combining both provided greater separation. Joint model recovery was significantly above chance for all three mechanisms (persistent-state, 46%, *p* = 8.5 × 10^−9^; controller, 44%, *p* = 1.1 × 10^−5^; teaching, 48%, *p* = 4.6 × 10^−11^, binomial test; **Supplementary Fig. S1**).

Recovery remained robust across broad parameter variations (**Supplementary Fig. S2**), and temporal-signature recovery improved with increasing trial number (**Supplementary Fig. S3**). Recovery also depended on behavioral accuracy, but remained measurable across the tested performance range (**Supplementary Fig. S4**). Finally, the predicted signatures were largely distinct across models: the persistent-state and controller models showed moderate similarity in their current-trial predictions (cosine similarity = 0.57), whereas the persistent-state and teaching models showed weaker similarity across trials (cosine similarity = 0.23; **Supplementary Fig. S5**). Together, these simulations show that the temporal structure of LC perturbation effects can distinguish the candidate computational mechanisms.

### Weak white-noise LC activation provides a temporally unbiased probe of perceptual behavior

We next applied this system-identification approach in monkeys performing a delayed match-to-sample orientation task. The excitatory opsin channelrhodopsin-2 (ChR2) was selectively expressed in noradrenergic LC neurons to permit optogenetic activation (**Fig. 3a**). Monkeys maintained fixation while two oriented sample stimuli were presented in opposite visual hemifields for 200 ms (**Fig. 3b**). Following a 200 ms delay, a test stimulus appeared at one of the two locations with equal probability. Animals reported whether its orientation matched that of the sample previously presented at the same location. Because the test location was unpredictable, the animals had to encode and retain the sample information at both locations.

**Figure 3.**
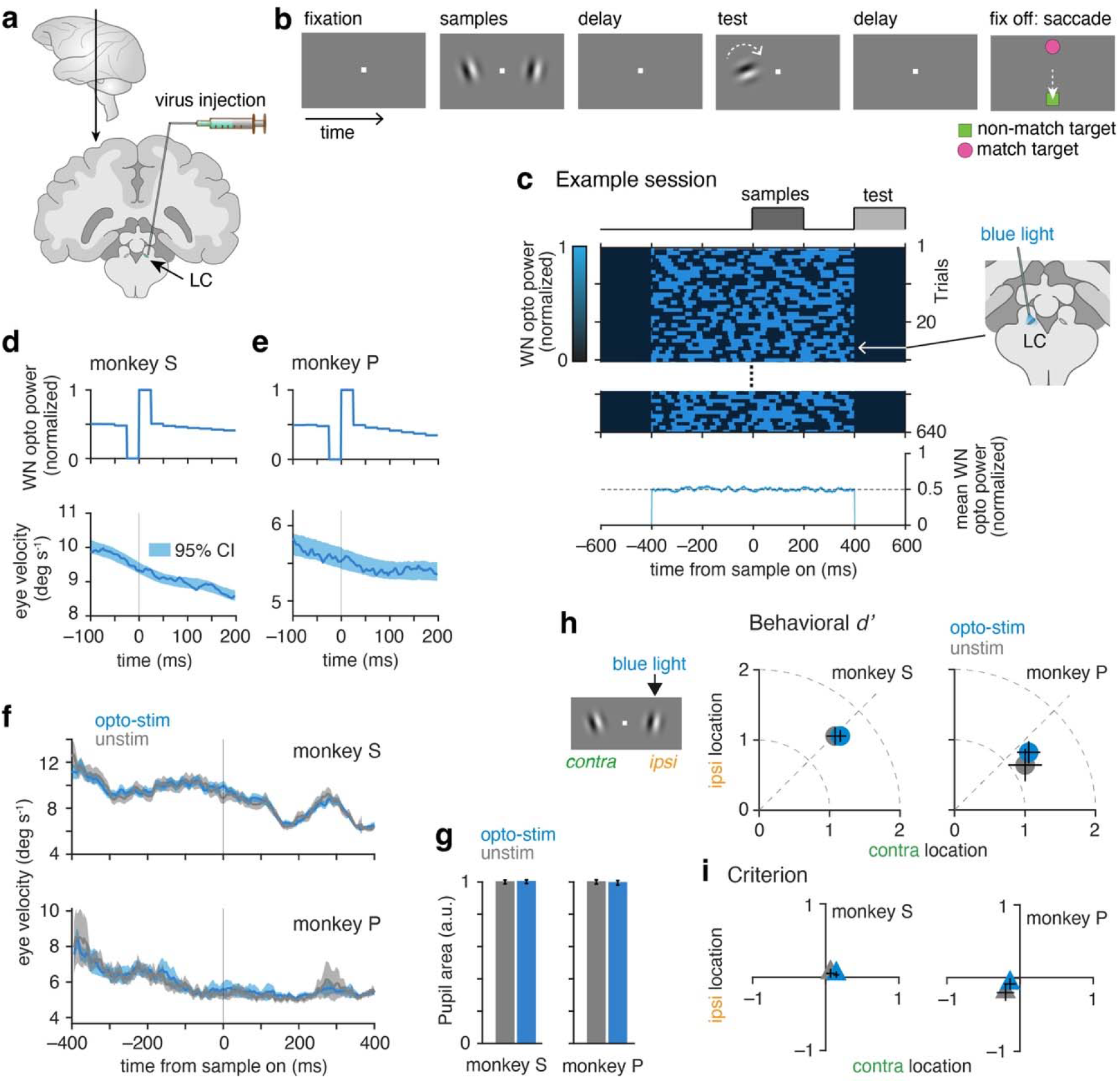
Experimental strategy for estimating the temporal contribution of LC activation to visual detection using weak white-noise optogenetic perturbations. (**a**) Schematic illustrating unilateral viral targeting to selectively express ChR2 in noradrenergic LC neurons in monkeys for optogenetic manipulation. (**b**) Delayed match-to-sample visual orientation-change detection task. Following central fixation, two Gabor sample stimuli were presented simultaneously in opposite visual hemifields for 200 ms. Following a 200 ms delay period, a test stimulus (200 ms) appeared at one of the two stimulus locations with equal probability. Monkeys reported whether the test stimulus matched (0.5 probability) the orientation of the sample at that location by making a saccade to match or nonmatch saccade-targets once the fixation-spot turns off (go-cue) after a brief delay. The positions of the two saccade-targets switched randomly across trials to dissociate motor-bias from perceptual decision criterion. (**c**) Weak white-noise (WN) optogenetic stimulation to the LC-NE neurons was delivered throughout an 800 ms period around the sample stimuli onset (–400 to 400 ms) in two-thirds of randomly interleaved trials. *Top*, trial-by-trial WN opto-stim patterns from an example session. Each row represents a trial and each column a time bin (25 ms). Blue and black indicate light-on and light-off states, respectively. Stimulation patterns were independently randomized across trials and time. *Bottom*, Average stimulation power across trials in the same example session remained constant throughout the stimulation epoch, demonstrating the absence of systematic temporal structure in the delivered perturbation. This design enabled reverse-correlation analysis to estimate the behavioral impulse-response function linking LC perturbations to perceptual performance. (**d, e**) Average absolute eye velocity aligned to onset of individual WN pulses for monkey S (d) and monkey P (e). *Top*, average change in normalized optogenetic power aligned to WN pulse onset. *Bottom*, mean eye velocity. Shading indicates the 95% interval of the null distribution obtained from 1,000 random temporal alignments. (**f**) Mean eye velocity aligned to sample onset for stimulated (blue) and unstimulated (gray) trials in monkey S (*top*) and monkey P (*bottom*). Shading indicates ±1 SEM across trials. (**g**) Mean pupil area during sample period for stimulated (blue) and unstimulated (gray) trials (normalized to unstimulated trials) for monkey S and monkey P. Error bars indicate ±1 SEM across sessions. (**h**) Behavioral sensitivity (*d’*) for monkey S (*left*) and monkey P (*right*). (**i**) Decision criterion for monkey S (*left*) and monkey P (*right*). In both panels, the horizontal and vertical axes represent contralateral and ipsilateral test locations, respectively; gray and blue symbols denote unstimulated and stimulated trials. Trials across all sessions were combined. Error bars denote 95% confidence interval. Weak WN stimulation produced no reliable changes in behavioral sensitivity or criterion.

To estimate the temporal contribution of LC activation without imposing a constrained temporal structure, we delivered weak binary WN stimulation from 400 ms before to 400 ms after sample onset on randomly interleaved stimulated trials (**Fig. 3c**), adapting an optogenetic reverse-correlation approach previously used to recover the temporal contribution of neural activity to behavior ^21^. Optical stimulation was independently randomized across 25 ms time bins and across trials. Individual trials therefore contained different temporal patterns of LC activation, whereas the average stimulation power across trials remained approximately constant throughout the stimulation interval. This allowed the timing of LC activation to vary independently of task events and provided the temporal variation required to estimate its contribution to behavior.

An important requirement of this approach was that the WN stimulation act as a weak probe rather than produce a gross change in behavioral or arousal state. We therefore first examined its oculomotor and pupil consequences. Aligning eye velocity to individual transitions in optical power revealed only modest pulse-associated changes in both monkeys (**Figs. 3d-e**), while eye-velocity profiles aligned to sample onset were closely overlapping between stimulated and unstimulated trials (**Fig. 3f**). Mean pupil area during the sample period was also similar between stimulation conditions in both animals (**Fig. 3g**). Thus, the stimulation did not produce an evident sustained change in these measures of oculomotor or pupil-linked state.

We next tested whether the WN perturbation produced any overall change in perceptual performance without accounting for stimulation timing. Behavioral sensitivity (*d’*) did not differ significantly between stimulated and unstimulated trials for both contralateral and ipsilateral test locations in each monkey (stimulated – unstimulated: Δ*d’*, monkey S, contralateral 0.053, *p* = 0.57, ipsilateral –0.001, *p* = 0.99; monkey P, contralateral 0.06, *p* = 0.75, ipsilateral 0.19, *p* = 0.31; permutation test; **Fig. 3h**). Decision criterion was likewise similar between stimulation conditions and test locations (Δcriterion, monkey S, contralateral 0.09, *p* = 0.07, ipsilateral –0.014, *p* = 0.8; monkey P, contralateral 0.02, *p* = 0.93, ipsilateral 0.13, *p* = 0.19; permutation test; **Fig. 3i**). Thus, weak WN LC activation did not produce any significant trial-averaged change in perceptual sensitivity or decision bias.

These results establish the regime required for system-identification analysis. The stimulation introduced temporally randomized variation in LC input without imposing an average temporal structure or producing a pronounced change in overall task performance. We could therefore ask how the precise timing and recent history of brief LC activation systematically influenced perceptual performance. The theoretical and simulation analyses above establish how the resulting within- and across-trial temporal weights can, in turn, distinguish the computational policies proposed in **Fig. 1**.

### Transient LC activation selectively contributes to contralateral perceptual performance within the ongoing trial

We next asked which of the predicted temporal signatures were expressed in perceptual behavior. Using the GLM framework validated in simulation, we estimated the contribution of WN LC activation as a function of its timing within the current trial and its history across preceding trials, separately for contralateral and ipsilateral detection in each monkey.

Although optical stimulation did not produce a clear overall change in trial-averaged task performance, a temporally localized within-trial contribution was evident in both monkeys (**Figs. 4a** and **4e**). For contralateral detection, GLM weights showed a narrow positive peak approximately 80 ms before sample onset (–83 ms, monkey S; –78 ms, monkey P). This temporal feature was absent for ipsilateral detection. A signal-detection analysis further showed that stimulation during the effective pre-sample window increased contralateral perceptual *d’* in both monkeys (monkey S, Δ*d’* = 0.222, 95% CI [0.128, 0.315]; monkey P, Δ*d’* = 0.206, 95% CI [0.039, 0.373]; **Figs. 4b, 4f, Supplementary Fig. S6**). These *d’* changes were significantly greater than those for ipsilateral detection (contra–ipsi difference, monkey S, 0.251, 95% CI [0.122, 0.380]; monkey P, 0.344, 95% CI [0.111, 0.577]), whereas no detectable changes in response criterion were observed. Thus, the behavioral consequence of LC optogenetic activation depended on both its timing and the location of the subsequently tested stimulus, consistent with a transient, spatially selective controller-like contribution to perceptual processing.

**Figure 4.**
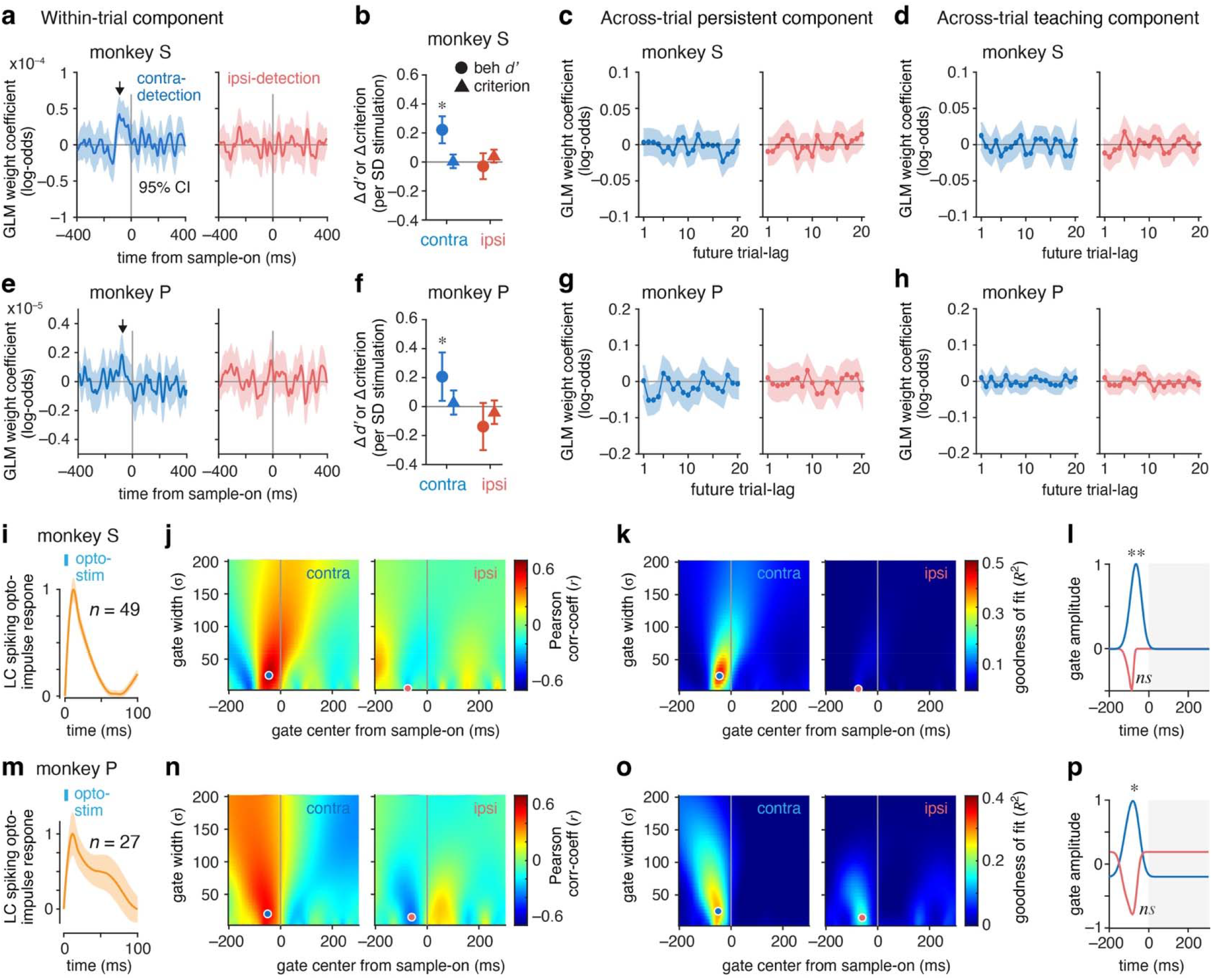
Temporal contributions of transient LC activation to perceptual performance within and across trials. (**a, e**) Within-trial contribution of white-noise LC activation to perceptual performance in monkey S (a) and monkey P (e). GLM coefficients are plotted as a function of activation time relative to sample onset, separately for contralateral (blue) and ipsilateral (red) detection. (**b, f**) Changes in behavioral *d’* and decision criterion associated with a one-standard-deviation increase in WN stimulation during the pre-sample window for contralateral and ipsilateral detections in monkey S (b) and monkey P (f). Error bars, 95% confidence intervals. Asterisks indicate effects whose 95% CIs exclude zero. (**c, g**) Across-trial persistent component, showing the contribution of LC activation on preceding trials to performance on subsequent trials. (**d, h**) Across-trial teaching component, showing the contribution of preceding LC activation, conditioned on signed behavioral outcome, to subsequent performance. In a, e, c-d, and g-h, coefficients are expressed as changes in log-odds of a correct response; lines show coefficient estimates and shading indicates 95% confidence interval. (**i, m**) Population LC spiking impulse response to optogenetic stimulation in monkey S (i) and monkey P (m). Shading indicates ±1 SEM across optically-activated LC units; *n* denotes the number of units. (**j, n**) Search for the timing of the LC controller. Candidate Gaussian temporal gates, defined by their center (μ) and width (σ) relative to sample onset, were forward-projected through the measured LC impulse response and compared with the observed optical-input-to-behavior kernel. Color indicates the Pearson correlation (*r*) between predicted and observed kernels for contralateral and ipsilateral detection. Circles mark the best-fitting gate parameters. (**k, o**) Goodness of fit of the same forward model across candidate gate centers and widths, quantified by zero-baseline *R*^2^. Circles mark the best-fitting parameters. (**l, p**) Best-fitting temporal gates for contralateral (blue) and ipsilateral (red) detection for monkey S (l) and monkey P (p), with amplitude determined by the corresponding GLM fit. Asterisks indicate significant fits (**, *p* < 0.01; *, *p* < 0.05) and *ns* indicates non-significant fits.

In contrast, we found no evidence for persistent or teaching effects across trials: across-trial weights showed no systematic decay across the 20 future-trial lags in either monkey or detection location (**Figs. 4c** and **4g**), contrary to the prediction of the persistent-state model. Similarly, LC activation conditioned on signed behavioral outcome did not systematically influence subsequent performance (**Figs. 4d** and **4h**), providing no evidence for the predicted outcome-dependent teaching component. Together, these analyses revealed a temporally restricted and spatially selective effect within the ongoing trial, while neither persistent LC-history nor outcome-dependent teaching-history predictors improved behavioral prediction on held-out trials relative to the within-trial controller model (monkey S, for contra, persistent, *p* = 0.87, teaching, *p* = 0.92, for ipsi, persistent, *p* = 0.70, teaching, *p* = 0.68; monkey P, for contra, persistent, *p* = 0.13, teaching, *p* = 0.99, for ipsi, persistent, *p* = 0.95, teaching, *p* = 0.58; paired permutation tests).

The absence of cross-trial contributions in the empirical data was unlikely to reflect substantially lower sensitivity of the GLM to history-dependent effects. The cross-trial kernels could be reliably recovered for behavioral effects comparable in magnitude to those required for within-trial kernel recovery (smallest tested stimulation-induced change in correct detection probability: within-trial controller, 0.33%; persistent-state history, 0.58%; teaching history, 0.38%; 95% confidence intervals of mean recovered GLM coefficients excluding zero, 2,000 trials; **Supplementary Fig. S7**).

We next asked whether the timing of the underlying LC spiking response could account for the observed within-trial behavioral effect. Because optical stimulation recruits LC spiking over tens of milliseconds, the timing of the optical-input kernel does not directly specify the timing of the corresponding LC activity. We therefore incorporated the population LC spiking impulse response to optical stimulation measured during neurophysiology sessions in each monkey (**Figs. 4i** and **4m**). Candidate transient Gaussian temporal gates were forward-projected through these measured LC impulse response dynamics. The resulting predicted optical-input kernels were compared with the observed WN-behavioral kernels across candidate gate centers and widths. In both monkeys, the contralateral fits identified a narrow pre-sample gate. Complementary correlation- and goodness-of-fit analyses converged on similar temporal parameters (**Figs. 4j-k, 4n-o)**. In monkey S, the best-fitting gate was centered 45 ms before sample onset with σ = 25 ms (Pearson correlation coefficient, *r* = 0.64, *p* = 0.001; *R*^2^ = 0.44, *p* = 0.004), whereas in monkey P it was centered 50 ms before sample onset with σ = 20 ms (correlation coefficient, *r* = 0.53, *p* = 0.018; *R*^2^ = 0.28, *p* = 0.034). The corresponding fitted temporal gates are shown in **Figs. 4l** and **4p**. Ipsilateral fits were not significant in either monkey (monkey S, µ = –75 ms, σ = 5 ms, correlation coefficient, *r* = –0.27, *p* = 0.81; *R*^2^ = 0.08, *p* = 0.88; monkey P, µ = –60 ms, σ = 15 ms, correlation coefficient, *r* = –0.38, *p* = 0.25; *R*^2^ = 0.18, *p* = 0.25).

Together, these results localize the behavioral contribution of LC activation to a brief period preceding sensory input. Its temporal and spatial specificity, together with the absence of persistent or outcome-dependent effects across trials, supports a transient controller-like mechanism rather than persistent-state or teaching mechanisms.

### The LC–behavior temporal kernel predicts contralateral perceptual performance on individual trials

We next asked whether the temporal structure identified by reverse correlation predicted behavioral outcome on individual trials. In a leave-one-out cross-validation, we projected the WN stimulation pattern from each held-out trial onto an LC-behavior kernel estimated independently from the remaining trials. The resulting projection quantified how strongly the timing of LC activation on that trial aligned with the temporal sensitivity profile identified by reverse correlation.

For contralateral detection trials, projection magnitude systematically predicted behavioral outcome in both animals. Binning trials into quartiles of projection strength revealed a progressive increase in detection performance with increasing projection value (monkey S: ρ = 0.07, *p* < 10^−4^; monkey P: ρ = 0.06, *p* = 0.04; Spearman correlation coefficient; **Figs. 5a, 5c** and **5g**). Trials with WN stimulation better aligned to the LC-behavior kernel were more likely to be correct. Receiver operating characteristic (ROC) analysis confirmed that projection values discriminated correct from error trials above chance. In monkey S, contralateral detection yielded an AUC of 0.55 (*p* < 10□^3^; permutation test), and in monkey P, an AUC of 0.54 (*p* = 0.038; **Figs. 5d** and **5h**). Discrimination was modest, but this was expected given the weak perturbation regime used to identify temporal sensitivity without producing a large trial-averaged behavioral effect. In contrast, identical analyses for ipsilateral detection showed no systematic relationship between projection and behavioral outcome. Projection quartiles did not exhibit a progressive trend (monkey S: ρ = ™0.01, *p* = 0.81; monkey P: ρ = 0.02, *p* = 0.26; **Figs. 5b, 5e** and **5i**), and classification performance did not differ from chance (monkey S: AUC = 0.49, *p* = 0.81; monkey P: AUC = 0.51, *p* = 0.26; **Figs. 5f** and **5j**).

**Figure 5.**
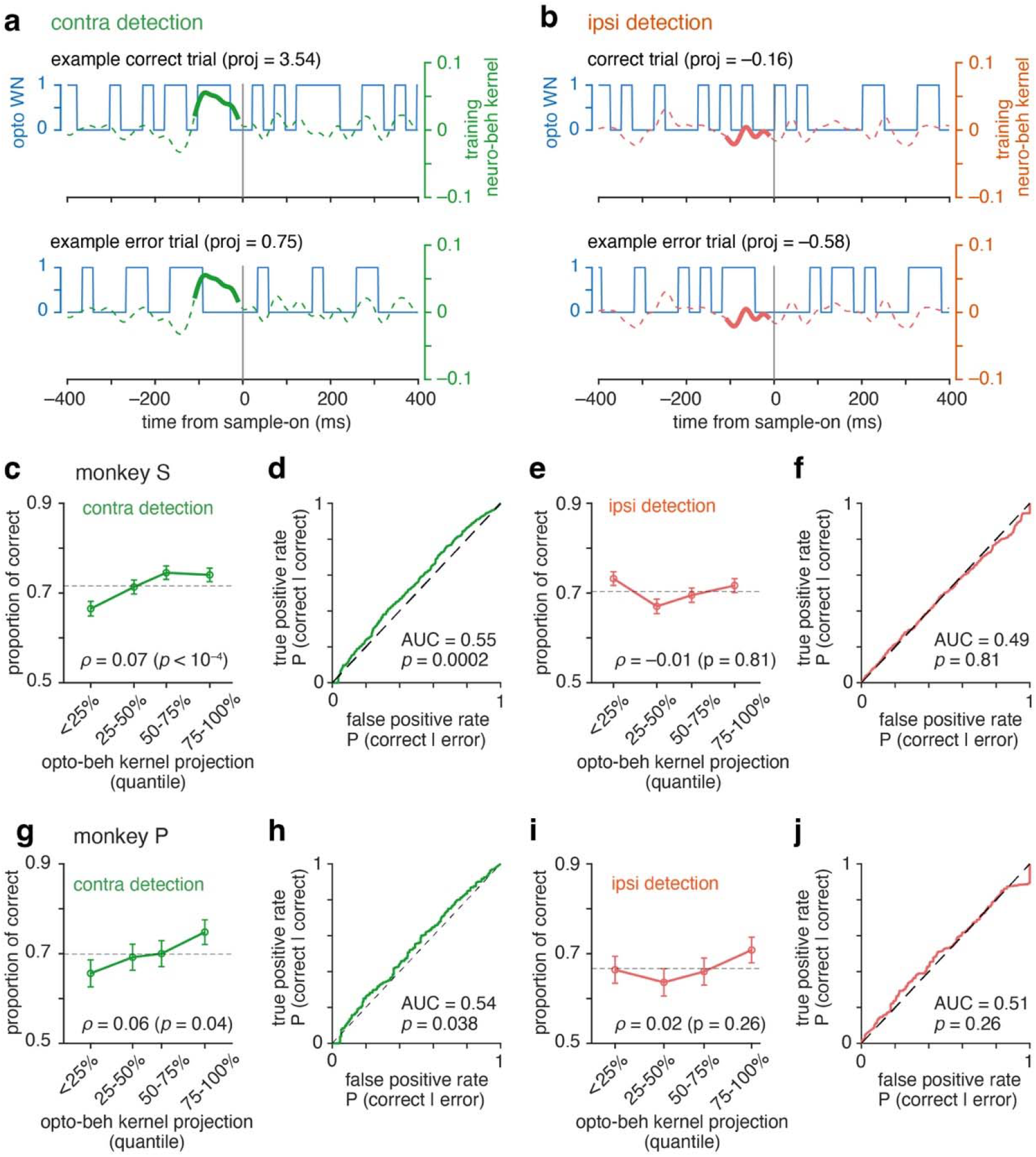
Temporally structured LC perturbations selectively predict contralateral detection on single trials. (**a, b**) Example trials illustrating projection of trial-specific WN optogenetic stimulation onto the cross-validated LC–behavior kernel. Blue traces show WN stimulation aligned to sample onset (vertical line). Green (a; contralateral) and red (b; ipsilateral) traces show the training kernel within the analysis window. Representative correct (top) and error (bottom) trials are shown with corresponding projection values. For contralateral trials (a), correct trials exhibited larger positive projections than error trials, reflecting stronger alignment between stimulation timing and the temporally defined behavioral sensitivity profile. For ipsilateral trials (b), projection values did not distinguish behavioral outcome. (**c, g**) Trial-level projection of white-noise optogenetic stimulation onto the LC-behavior kernel was binned into quartiles (cross-validated; leave-one-out) for contralateral detection in monkey S (c) and monkey P (g). Higher projection values were associated with increased probability of correct detection. Dashed lines indicate mean performance across trials. Spearman correlation (ρ) and permutation-based *p*-values are shown. (**d, h**) Receiver operating characteristic (ROC) analysis of cross-validated projection values for contralateral detection in monkey S (d) and monkey P (h). Projection significantly discriminated correct from incorrect trials (monkey S: AUC = 0.55, *p* <10□^3^; monkey P: AUC = 0.54, *p* = 0.038; permutation test). (**e, i**) Identical analysis for ipsilateral detection in monkey S (e) and monkey P (i). Projection did not systematically predict behavioral outcome, and no monotonic relationship between projection quartile and performance was observed (Spearman ρ near zero; permutation *p* > 0.25). (**f, j**) ROC analysis for ipsilateral detection. Classification performance did not differ from chance (monkey S: AUC = 0.49, *p* = 0.81; monkey P: AUC = 0.51, *p* = 0.26; permutation test).

Together, these analyses show that the timing of LC activation carried predictive information about perceptual outcome on individual trials, selectively for contralateral detection. The same spatial specificity observed in the temporal kernels therefore generalized to held-out trials, arguing against a nonspecific relationship between overall stimulation and behavioral performance.

## DISCUSSION

Neuromodulatory systems are commonly assigned broad functions such as arousal, gain regulation, learning or behavioral-state control, but these descriptions do not specify the role that a transient neuromodulatory event plays in an ongoing computation. Here, temporally randomized LC activation revealed a brief and spatially selective causal contribution to perceptual behavior. LC activation improved contralateral detection on the current trial when delivered within a narrow period approximately 80 ms before sensory input, but produced no detectable persistent or outcome-dependent influence across subsequent trials. Among the computational policies considered here, this combination most closely matched a transient controller architecture rather than a persistent state or teaching mechanism. Thus, the behavioral consequence of transient LC activity was determined not simply by whether LC was activated, but by when it was activated relative to the ongoing computation and where the behaviorally relevant sensory information subsequently appeared.

### Transient LC operation and persistent downstream consequences

These findings suggest an important distinction between the computational role played by a phasic LC event and the duration of its downstream consequence. A transient control signal need not produce only transient effects. If phasic LC activity modulates rapidly evolving sensory processing, as in the present task, its behavioral consequence may be brief. In other contexts, a computation transiently controlled by LC may itself generate longer-lasting output, without requiring the LC-dependent control signal to persist. Consistent with this possibility, brief LC stimulation can be followed by sustained changes in downstream neural activity, including prefrontal activity and distributed cortical network configuration ^11,12^. Such observations demonstrate persistence in a downstream consequence, but do not establish that the phasic LC signal itself implements a persistent state. In the present task, where brief LC activation could have displaced a state carried across trials, we detected no such carryover.

This distinction also clarifies the relationship between the controller framework and adaptive-gain accounts of LC function ^1^. Adaptive gain specifies a potential consequence of noradrenergic modulation, but does not by itself determine its temporal architecture. Gain modulation could arise through a persistent change in operating conditions or be deployed transiently when LC activity coincides with a computation susceptible to modulation. This distinction may also help separate the computational roles of sustained and phasic LC activity, which can produce different patterns of cortical engagement ^8,11^. The present results favor the latter architecture for the perceptual effect examined here.

### Transient LC control during learning

A similar distinction applies to learning. The animals in the present study performed a well-learned perceptual task with stable stimulus-response contingencies, rather than acquiring or substantially updating a behavioral policy. Accordingly, the absence of an outcome-dependent across-trial effect does not exclude a role for phasic LC in learning when behavioral policies are actively being acquired or updated. Learning-related LC activity and causal effects of LC stimulation have been observed during acquisition, unexpected outcomes and changing contingencies ^5,17-19^. Our teaching model tested the more specific hypothesis that LC activation interacts with signed trial outcome to generate an update expressed in subsequent behavior, for which we found no evidence under the present task conditions.

Learning nevertheless provides a clear way in which transient control could generate consequences expressed on subsequent trials. Learning depends on temporally structured computations, including outcome evaluation and the updating of neural representations or action policies. These processes therefore provide potential targets for transient LC control. If phasic LC coincides with a period of plasticity, it could transiently regulate the efficacy of the ongoing update, while information specifying what is learned could be supplied by the downstream learning process ^17^. The resulting update could be stored by that circuit and influence subsequent behavior after the LC signal has ended. This differs from a persistent-state mechanism because the LC-dependent control signal is not retained, and from a teaching mechanism because LC does not specify the content or direction of the update. It predicts that during active learning, the effect of LC perturbation should depend on its timing relative to the learning computation, which could be tested using temporally randomized LC perturbations during acquisition or changing contingencies.

### Temporal and spatial specificity of LC control

The controller interpretation provides a natural account of the temporal and spatial specificity observed here. After incorporating the measured LC spiking response to optical stimulation, a narrow period of behaviorally effective LC activity shortly before sensory input was identified in both monkeys. Timing-dependent modulation of sensory cortical responses by phasic LC activation has also been observed previously ^8,16^. Spatially, unilateral LC activation affected detection selectively in the contralateral visual field, consistent with our previous finding that LC activation selectively improves contralateral perceptual sensitivity ^7^. Together, these observations suggest that the consequence of a phasic LC event depends on the computation susceptible to modulation at that moment rather than being specified by LC activation alone.

More generally, the diversity of functions attributed to LC need not imply that phasic LC implements a different computational policy in each context. A parsimonious possibility is that phasic LC has a characteristic controller-like operation, while the identity and dynamics of the downstream computation determine what is controlled and how long the resulting consequence lasts. Distinguishing the temporal structure of the neuromodulatory operation from that of its downstream consequences may therefore provide a general framework for understanding how transient neuromodulatory signals influence behavior across timescales.

## METHODS

### Animals and surgery

Two adult male rhesus monkeys (*Macaca mulatta*, 9 and 13 kg) participated in the experiments. Both animals had previously been implanted with titanium head posts and recording chambers unilaterally targeting the LC with guidance from structural MRI (right LC in monkey S; left LC in monkey P). The same animals were used in a previous study ^7^. The viral injections, targeting procedures and expression of channelrhodopsin in LC noradrenergic neurons were performed as described previously ^7^. No additional viral injections were performed for the present study. The same two monkeys were used in previous studies that described different findings on neuronal responses in superior colliculus ^23^ and LC ^7^. All experimental procedures were approved by the Institutional Animal Care and Use Committee of the University of Chicago and followed the U.S. National Institutes of Health guidelines.

### Behavioral task

Monkeys performed a visual match-to-sample orientation-discrimination task that has been described previously ^23^. Animals sat in a primate chair facing a calibrated CRT display (1024 × 768 pixels, 100 Hz refresh rate) at a viewing distance of 57 cm in a darkened room. Binocular eye position and pupil area were recorded at 500 Hz (EyeLink 1000, SR Research). Each trial began when the animal acquired fixation within 1.5° of a central fixation point. After 400-800 ms of fixation, two achromatic Gabor sample stimuli appeared for 200 ms, one in each visual hemifield. Following a delay of 200 ms, a 200 ms test Gabor stimulus appeared with equal probability at either sample location. The spatial phases of the sample and test Gabor stimuli were fixed (0°, odd symmetric). On half of trials, the test orientation differed from the corresponding sample orientation (non-match trial); on the remaining trials, the orientations were identical (match trial). At 100-150 ms after test offset, two saccade targets appeared (non-match, green square; match, magenta circle; 0.30°-0.55°), followed 150-200 ms later by a go-signal (fixation spot offset) instructing the animal to report match or non-match by saccade. Saccade-target locations were randomized across trials to dissociate the perceptual decision from a fixed motor response. Sample orientations at each location were independently randomized from 0° to 175° in 5° increments. Stimulus positions and sizes were selected for each animal as described previously and remained fixed within individual recording sessions. Critically, the two test locations occurred with equal probability and correct responses at the two locations were associated with equal reward magnitude. Thus, unlike conditions designed to manipulate spatial attention through reward, the animals had no differential reward incentive to prioritize either location.

### Viral expression of channelrhodopsin in LC noradrenergic neurons

Viral targeting and expression of channelrhodopsin in LC noradrenergic neurons were previously verified histologically in these same animals ^7^. Briefly, a dopamine-β-hydroxylase promoter-driven Cre vector (AAV9-DBH-Cre-P2A-mCherry-SV40pA) was combined with a Cre-dependent channelrhodopsin vector (AAV5-EF1α-DIO-hChR2(H134R)-EYFP) and injected unilaterally in and around the electrophysiologically identified LC. The injections and subsequent expression period preceded the experiments reported here; no additional viral manipulation was performed for the present study.

### LC recordings

LC activity was recorded to measure the LC spike impulse response to optogenetic stimulation, using linear multielectrode arrays incorporating optical fibers (16-channel V-probes with two optical-fiber channels; Plexon). These recording sessions were different from the optogenetic WN stimulation sessions. Extracellular signals were amplified, band-pass filtered (250-7500 Hz) and sampled at 30 kHz (Cerebus, Blackrock Microsystems). Spikes were sorted offline using principal-component and waveform features (Offline Sorter, Plexon). Well-isolated clusters were classified as single units based on waveform separation and cluster isolation, as described previously^7^. LC recording sites were identified from stereotaxic location and characteristic electrophysiological response properties, including responses to brief arousing auditory stimuli, together with anatomical landmarks encountered along the electrode trajectory.

### White-noise optogenetic stimulation of LC

Optical stimulation was delivered to the transduced LC using a 457-nm laser (Laserglow Technologies), with laser power controlled by a Pockels cell (ConOptics) through the behavioral-control software. Light was delivered through an optical fiber (100 µm core cross-section, NA, 0.66, Doric Lenses) inserted into the LC. On two-thirds of trials, LC was stimulated with an 800 ms binary white-noise (WN) sequence extending from 400 ms before to 400 ms after sample onset. The WN sequence consisted of 25 ms bins in which the light stimulus was either on or off. The phase of the WN bin sequence relative to sample onset was varied across trials, so that pooled stimulation transitions sampled task time at finer than 25 ms resolution. A weak optical stimulation intensity (< 10 mW at the fiber tip) was used, substantially lower than the 50 mW intensity used to produce behavioral effects in our previous study ^7^. The remaining trials were randomly interleaved unstimulated trials. This temporally randomized stimulation allowed the contribution of LC activation to behavior to be estimated across task time without restricting stimulation to a predefined behavioral epoch.

## Data analysis

### Computational models and predicted temporal signatures

To distinguish competing roles of transient LC activity, we implemented three computational models that received the same temporally patterned LC input but differed in how that input influenced behavior. The persistent-state model treated LC activation as a perturbation of a latent operating state that could influence multiple task computations and persist beyond the initiating LC event. The controller model treated LC activation as a transient control input whose efficacy depended on a computation-specific temporal gate. The teaching model treated LC activation as part of an outcome-dependent update that affected behavior only on subsequent trials. Model dynamics were integrated at 5 ms resolution. Temporal behavioral kernels were not imposed on the models but were derived from their governing dynamics and subsequently recovered from simulated behavior using the analyses described below.

#### Persistent-state model

In the persistent-state model, LC input perturbed a latent operating state whose dynamics were slower than the input itself. The LC-driven state decayed with a time constant that depended on the background state and ranged from 50 to 300 ms. The background operating state included slow trial-to-trial fluctuations and recent-performance dependence, together with trial-locked temporal expectation and within-trial endogenous variability. These components provided a dynamically varying background state but did not generate the diagnostic LC effect. The total state was,

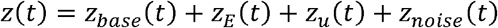

where *z*_base_(*t*) is the operating-state baseline carried across trials, *z*_E_(*t*) is an expectation-related background state and *z*_u_(*t*) the component driven causally by LC WN input, and *z*_noise_(*t*) represents endogenous within-trial fluctuations. The LC-driven state followed leaky dynamics, and did not directly enter the behavioral decision. Instead, the common operating state modulated sensory encoding of the sample and test stimuli, retention of the sample representation during the delay, decision criterion and lapse probability. The LC-driven component remaining at the end of the stimulation interval was partially written into the operating-state baseline of the following trial, with a retention factor of 0.8 across subsequent trials. Thus, a brief LC perturbation could influence several computations within the current trial and could also produce a decaying effect on future trials.

#### Controller model

The controller model assumed that LC input *u*(*t*) influenced behavior only when it coincided with a computation-specific temporal gate *G*(*t*),

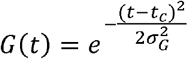

where *t*_c_ is the center of the susceptible temporal window and *σ*_G_ = 35 ms is its temporal width. Candidate gate centers were 0 and 100 ms relative to sample onset.

Gated LC input drove a transient controller state *c*(*t*) with a time constant *τ*_c_ = 25 ms,

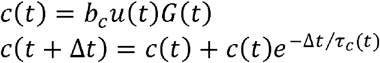

with an LC-to-controller input gain, *b*_c_ = 0.2. The controller signal modulated the currently active sensory representation with a gain,

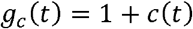

but did not alter memory retention, decision criterion or lapse probability and was not retained across trial boundaries. Consequently, the model predicted a temporally localized effect on current-trial behavior, which depended on the temporal overlap between LC input and the computation-specific gate. This effect shifted with the timing of the temporal gate but did not show across-trial carryover.

#### Teaching model

In the teaching model, LC input did not alter the perceptual computation generating the current choice. Instead, trial-level LC drive *u*_n_ interacted with signed behavioral outcome feedback *δ*_n_ (+1 for a correct response and –1 for an incorrect response) after the choice, to generate a teaching signal,

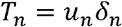

The teaching signal updated a retained policy weight,

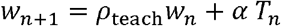

where, *ρ*_teach_ = e^−1/3^ is the retention factor and *α* = 0.2 is the learning rate. The policy weight contributed to behavioral log-odds on subsequent trials. Because the update occurred after the current choice, the teaching model predicted no direct within-trial effect of LC activation but an outcome-dependent influence on future behavior.

#### Perceptual readout

The persistent-state and controller models shared the same perceptual readout. LC-dependent modulation altered the strength of the sample and test representations and thereby changed the precision of their comparison, without changing the physical orientation difference. Effective comparison noise, *σ*_eff_ was determined by the relative strength of the sample and test representations as,

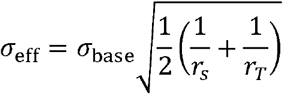

where, *r*_*S*_ and *r*_*T*_, are the strengths of the sample and test representations relative to their unmodulated values, respectively, and *σ*_base_ = 14° is the baseline comparison noise. For nonmatch trials, the physical orientation difference between the sample and test stimuli was Δ*θ* = 20°, whereas for match trials Δ*θ* = 0°. An orientation difference was perceived as change when the magnitude of its noisy internal estimate exceeded the decision criterion *C* = 10°,

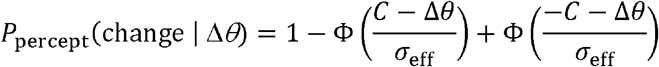

where, Φ(•) is standard normal cumulative distribution function. A 5% lapse probability (λ =0.05) was incorporated at the response stage,

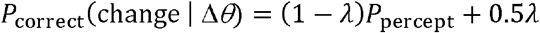

Match and nonmatch trials were generated with equal probability.

#### Prediction of within- and across-trial signatures

The theoretical temporal signatures in **Figs. 1e-f** were obtained directly from the deterministic model dynamics before generating stochastic behavioral choices. For the within-trial prediction, LC input was applied to one 25 ms stimulation bin at a time, and the resulting probability of a correct response was compared with the corresponding no-LC input condition. The behavioral contribution was expressed as

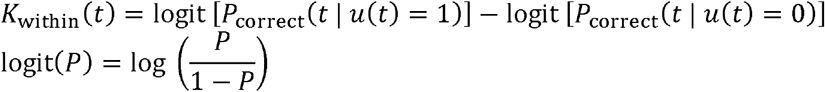

Thus, *K*_within_(*t*) expresses the behavioral consequence of an isolated LC perturbation at time *t* in log-odds units. The persistent-state model produced a broad current-trial kernel because an LC-induced state displacement continued after the perturbation itself. The controller produced a temporally restricted kernel determined by the controller gate. Because the teaching update occurred only after trial outcome, the teaching model predicted no current-trial kernel.

Cross-trial predictions were calculated by propagating the model-specific consequence of an LC event across subsequent trials. For the persistent-state model, LC perturbation-induced displacement of the operating-state baseline decayed across trials with the across-trial retention factor *ρ*_state_ = 0.8. For the teaching model, the policy-weight update decayed across trials according to *ρ*_teach_, producing an outcome-dependent across-trial behavioral effect. The controller carried no state across trial boundaries and therefore predicted zero across-trial influence. The models could therefore be distinguished jointly: persistent-state and controller mechanisms could both affect the current trial, but only persistence carried across trials; persistent-state and teaching mechanisms could both affect subsequent trials, but only teaching depended on trial outcome.

### Behavioral simulation and recovery of within- and across-trial kernels

#### Stochastic behavioral simulation

To determine whether the predicted temporal signatures could be recovered from stochastic behavioral observations, we generated simulated datasets from each computational model. LC input consisted of an independently randomized binary WN sequence spanning 400 ms before to 400 ms after sample onset. WN states were independently drawn in 25 ms bins and represented by five consecutive 5 ms samples for integration of the model dynamics. Thus, model dynamics were evaluated at 5 ms resolution, whereas the statistically independent perturbation variables used for kernel recovery corresponded to the 32 physical 25 ms WN bins. The same WN realizations and task variables were supplied to each model so that differences in simulated behavior arose from the model-specific transformation of LC input.

On each trial, sample orientation was drawn uniformly from 0° to 180°. Match and nonmatch trials occurred with equal probability; on nonmatch trials, test orientation differed from the sample by 20°, with the sign of the difference assigned randomly. For the teaching model, signed outcome feedback *δ*_n_ was defined as +1 for a correct response or –1 for an incorrect response and combined with trial-level LC drive to generate the teaching signal *T*_n_ = *u*_n_*δ*_n_, as defined above. Each model generated an exact trial-wise probability of correct performance, *P*_correct,n_, from which a binary correct or incorrect behavioral outcome was sampled. Thus, the simulated behavioral datasets incorporated finite binary response-sampling variability in addition to the model dynamics.

#### Recovery of within-trial kernels

Within-trial kernels were recovered directly from the independently randomized WN states without imposing the theoretically predicted temporal profiles. For each trial, the 32 WN states were entered simultaneously as predictors of binary behavioral outcome in a logistic generalized linear model (GLM),

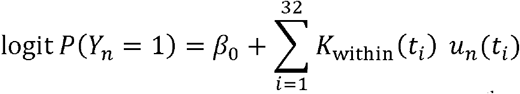

where, *Y*_*n*_ is behavioral outcome and *u*_*n*_(*t*_i_) is the WN state in the *i*^th^ 25 ms bin. The fitted coefficients *K*_within_(*t*_*i*_) constituted the recovered within-trial kernel in log-odds units. Thus, the contribution of each independently randomized WN bin was estimated while controlling for stimulation in all other bins. The GLMs were fit without regularization (*λ* = 0). In parallel, a population kernel was estimated using the same design but fitting the exact model-generated probabilities, *P*_correct,n_, rather than sampled binary outcomes. The population kernel therefore represented the expected GLM-recoverable signature in the absence of binary response-sampling noise.

#### Recovery of across-trial kernels

Across-trial kernels were recovered over the preceding 20 trials using history variables corresponding to the quantity propagated across trials by each hypothesis. For the persistent-state model, the WN sequence on each trial was propagated through the LC-driven component of the persistent-state dynamics described above. Only the WN-driven component was retained for this history predictor; performance history and endogenous state noise were excluded. The value of *z*_WN,n_(*t*) at the end of the WN stimulation interval was retained as the trial level persistent state drive, *s*_*n*_, and its across-trial mean was removed. For current trial *n*, the persistent-history vector was,

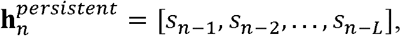

where *L* denotes the maximum modeled trial lag (*L* = 20). The lagged predictors were entered simultaneously into the logistic GLM

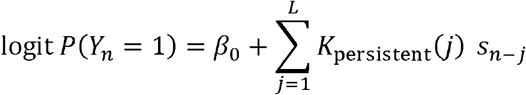

The fitted coefficients *K*_persistent_(*j*) constituted the recovered persistent-state across-trial kernel and quantified the association between the WN-induced persistent-state drive *j* trials before the current trial and performance on current trial *n*.

For the teaching model, the relevant history variable was the interaction between trial-level LC drive and signed outcome feedback. For current trial *n*, the teaching-history vector was

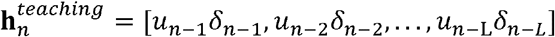

where *u*_n-j_ and *δ*_n-j_ are the LC drive and signed feedback respectively, from the trial *j* before the current trial. The lagged LC-by-feedback predictors were entered simultaneously into the logistic GLM,

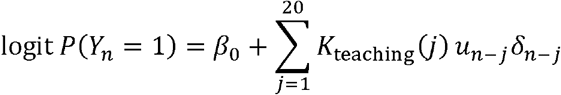

The fitted coefficients *K*_teaching_(*j*) constituted the recovered teaching across-trial kernel and quantified the outcome-dependent influence of preceding LC events on current-trial behavior. Thus, the two analyses tested distinct across-trial signatures: an outcome-independent WN-induced persistent-state drive for the persistent-state model and an outcome-dependent LC-by-feedback interaction for the teaching model. The controller retained no state across trial boundaries and therefore predicted no systematic across-trial kernel. The across-trial GLMs were fit without regularization (*λ* = 0). Corresponding population kernels were obtained by applying the same analyses to the exact model-generated probabilities of correct performance rather than to sampled binary outcomes, providing the expected recoverable across-trial signatures in the absence of binary response-sampling variability.

#### Sensitivity of across-trial and within-trial kernel recovery

To quantify the sensitivity of the GLM to cross-trial behavioral effects, we systematically varied the strength of the WN-dependent state update in the persistent-state model or the learning rate in the teaching model, holding all other model parameters fixed. For each strength, we generated 200 independent simulated datasets of 2,000 trials and recovered within-trial (32 WN bins) and across-trial (20 preceding trials) kernels using unregularized logistic GLMs, as described above.

The generating behavioral effect was quantified as the absolute difference in mean next-trial P(correct) between matched stimulation and no-stimulation conditions on the preceding trial. These counterfactual comparisons preserved preceding-trial feedback, earlier trial history, and current-trial inputs. Generating effects were averaged across three independent simulations and expressed in percentage points. Recovered GLM coefficients were converted to changes in predicted P(correct) for a one-standard-deviation increase in the corresponding predictor. For each generating effect strength, we calculated the mean recovered coefficient and its 95% confidence interval across simulations. We defined the recovery threshold as the smallest tested generating behavioral effect for which the lower 95% confidence bound of the mean recovered coefficient for trial-lag 1 exceeded zero.

Within-trial sensitivity was assessed separately by varying the strength of the controller mechanism and measuring the corresponding change in current trial probability of correct detection between matched stimulated and unstimulated conditions, using the same kernel-recovery and confidence-interval criteria.

#### Held-out model identification

Kernel recovery and model identification were performed as separate analyses. The preceding GLMs estimated temporal kernels without imposing their theoretically predicted shapes. Model identification instead tested whether the model-specific temporal signatures were sufficient to identify the computational mechanism that generated the behavior. For each candidate model, the within-trial WN sequence was projected onto that model’s theoretical within-trial kernel to generate a single current-trial predictor. For the persistent-state model, preceding-trial LC-induced state drive was similarly projected onto the theoretical persistent-state across-trial kernel; for the teaching model, the LC-by-feedback history was projected onto the theoretical teaching across-trial kernel. Because the controller predicted no across-trial persistence, its across-trial-only model contained no history predictor.

Simulated trials were randomly divided into 80% training and 20% held-out test sets. Candidate models were fit exclusively to the training trials without regularization, and predictive performance was evaluated on held-out binary behavioral outcomes using mean Bernoulli log likelihood. Model identification was evaluated separately using within-trial information, across-trial information, and their combination. The candidate model with the highest held-out log likelihood was designated the winner. Candidate implementations with different temporal gates were grouped into a common controller computational family for summary analyses, and recovery probability was calculated across independently simulated datasets as the proportion in which the generating computational family was correctly identified.

### Experimental measures of behavioral and physiological effects

#### Behavioral performance

Behavioral sensitivity (*d’*) and perceptual decision criteria (*C*) were calculated separately for stimulated and unstimulated trials and for each stimulus location (contralateral or ipsilateral to the stimulated LC) using one-dimensional signal detection theory ^24,25^. Hit rate (*H*) was defined as the proportion of nonmatch trials reported as nonmatch, and false-alarm rate (*F*) as the proportion of match trials reported as nonmatch. Sensitivity and criterion were calculated as

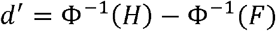

And

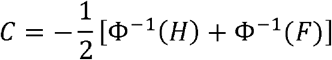

where Φ^−1^ is the inverse normal cumulative distribution function.

Behavioral analyses included 10,080 trials from 12 sessions for monkey S and 3,000 trials from 5 sessions for monkey P. For trial-averaged comparisons (**Figs. 3h-i**), response counts were pooled across sessions separately for each stimulation condition and test location. Statistical significance was assessed using 1,000 session-level paired permutations. In each permutation, stimulated and unstimulated labels were randomly swapped within each session, with all response counts for each condition kept together. Counts were then pooled across sessions, and stimulated-minus-unstimulated differences in d′ and criterion were recalculated. Two-sided P values were obtained from the resulting permutation distributions.

#### Pupil area

Pupil area was measured binocularly at 500 Hz using an infrared eye-tracking system (EyeLink 1000, SR Research). Pupil traces were aligned to sample onset and analyzed separately for each eye. For each recording session, mean pupil area was calculated for stimulated and unstimulated trials over a 250 ms period (0 to 250 ms after sample onset), pooling trials across contralateral and ipsilateral test trials. Session-level values were then normalized by the mean pupil area on unstimulated trials for the corresponding eye. Mean normalized pupil area and its SEM across sessions were calculated separately for stimulated and unstimulated trials.

#### Eye velocity

Eye position was measured binocularly at 500 Hz and aligned to sample onset. Horizontal and vertical eye-position traces were first smoothed using a five-sample moving average (10 ms), differentiated at 2 ms resolution to obtain horizontal and vertical eye velocity, and smoothed again using a five-sample moving average. Absolute two-dimensional eye velocity was calculated as

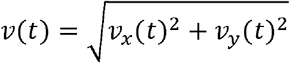

Eye-velocity traces were analyzed from 400 ms before to 400 ms after sample onset and averaged across trials separately for stimulated and unstimulated conditions. The mean and SEM of sample-aligned traces across trials were calculated at each time point (**Fig. 3f**).

To test whether individual WN stimulation events were associated with time-locked changes in eye movements, absolute eye velocity was additionally aligned to transitions in the WN stimulation sequence. A null distribution was generated using 1000 random temporal alignments, preserving the number of events in each iteration. The mean eye velocity trace was calculated for each random alignment, and the 95% confidence interval was estimated from this null distribution.

### Experimental analysis of within- and across-trial temporal contributions of LC probed by optogenetic white-noise

Reverse-correlation approaches have previously been used to estimate behavioral weighting functions from temporally or spatially varying sensory input and from optogenetic WN perturbations ^20-22^. We quantified the temporal contribution of LC activation to perceptual performance using logistic GLMs relating the optogenetic WN sequence to trial-by-trial behavioral outcome. Analyses were performed separately for trials in which the test stimulus appeared contralateral or ipsilateral to the stimulated LC. The empirical WN sequence was represented in 5 ms bins from 400 ms before to 400 ms after sample onset.

#### Within-trial LC-behavior (correct versus error) kernel

The within-trial kernel was estimated from stimulated trials using ridge-regularized logistic regression,

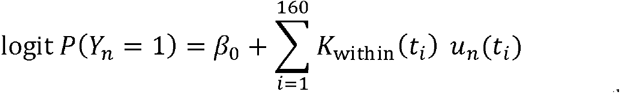

where *Y*_n_ denotes behavioral outcome and *u*_n_(*t*_i_) is the binary WN state in the *i*^th^5 ms bin. The fitted coefficients

*K*_within_(*t*_i_) quantified the contribution of LC stimulation at each time relative to sample onset in log-odds units. Ridge regularization was used while fitting. The regularization parameter *λ* was selected by 10-fold cross-validation from 37 logarithmically spaced values between 10^−6^ and 10^6^, and was then held fixed for the descriptive fit and bootstrap analysis. Current-trial WN predictors were retained in their native binary scale. Confidence intervals (95%) were estimated from 1000 bootstrap resamples of trials using the fixed *λ*.

#### Within-trial LC-behavior (d’ and criterion) kernel

The temporal effects of LC stimulation on perceptual *d’* and decision criterion were estimated from stimulated trials using ridge-regularized probit regression,

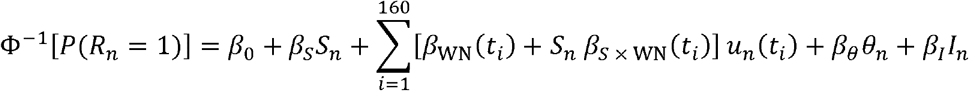

where, Φ^−1^ is the inverse standard normal cumulative distribution function, *β*_0_, *β*_*S*_, *β*_WN_, *β*_S×WN_, *β*_θ_, and *β*_*I*_ are fitted regression coefficients, *R*_n_ is the monkey’s report (0 = match, 1 = nonmatch), *S*_n_ is test stimulus identity (0 = match, 1 = nonmatch), *u*_n_(*t*_i_) is the binary WN state in the *i*^th^ 5 ms bin. *θ* _n_ and *I*_n_ are sample orientation and within-session trial number, respectively. The model was fitted separately for each stimulus location. Ridge regularization was applied to the WN and interaction coefficients, with λ = 1000 held fixed for fitting and bootstrap analysis. Under the equal-variance signal-detection model, the temporal kernels for perceptual *d’* and criterion were calculated as,

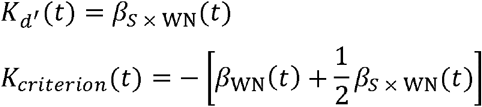

where, *β*_WN_(*t*) is the WN coefficient and *β*_S×WN_(*t*) the stimulus identity × WN interaction coefficient. Confidence intervals (95%) of temporal-kernels were estimated from 1000 trial-level bootstrap samples.

To quantify stimulation-induced changes in *d’* and criterion (**Figs. 4b** and **4f**), we fitted a separate probit GLM incorporating mean WN stimulation within the 120 ms window centered on the pre-sample peak of the within-trial LC-behavioral kernel (monkey S, –120 to 0 ms; monkey P, –140 to –20 ms) and outside this window, together with stimulus identity, location, their interactions with stimulation, and the same task variables. Effects were expressed per one-standard-deviation increase in within-window WN stimulation. Differences between stimulation-induced effects at contralateral and ipsilateral locations were estimated directly, with 95% confidence intervals calculated from the fitted coefficient covariance matrix.

#### Persistent across-trial analysis

To test whether LC activation produced an outcome-independent influence on subsequent behavior, WN stimulation on each trial was summarized as the mean WN state across the stimulation interval. This trial-level LC drive was standardized across eligible stimulated trials, with unstimulated trials assigned zero drive. For current trial *n*, the preceding 20 trial-level LC drives were entered simultaneously into a ridge-logistic GLM together with the current-trial WN sequence and nuisance covariates,

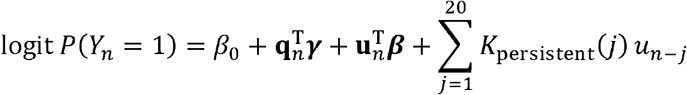

where **u**_n_ denotes the 160 element current-trial WN sequence, *β* its corresponding vector of temporal coefficients, and **q**_n_ contains experimental covariates (match/nonmatch trial type, sample orientation and within-session trial number), with corresponding coefficients *γ* ; **T** denotes transpose. These covariates were included to account for trial-to-trial variation in task conditions and slow within-session changes in performance that could otherwise contribute to apparent history effects. The coefficients *K*_persistent_(*j*) constituted the empirical persistent across-trial kernel.

#### Teaching across-trial analysis

To test whether preceding LC activation interacted with behavioral outcome to influence subsequent behavior, correct and incorrect responses were coded as *δ*_n_ = +1 and –1, respectively, and trial-level LC drive was multiplied by this signed outcome feedback. The corresponding history model was,

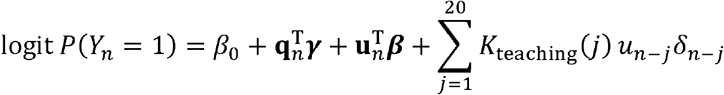

The coefficients *K*_teaching_(*j*) constituted the empirical outcome-dependent teaching signal across-trial kernel.

For both across-trial analyses, histories were constructed from the complete chronological trial sequence before separating current trials by test location. Histories did not cross recording-session boundaries and were restarted following discontinuities in block progression. Ridge penalties for the descriptive across-trial kernels were selected by 10-fold cross-validation and held fixed during 1000 bootstrap resamples of eligible trials pooled across sessions and blocks, and 95% confidence intervals were estimated.

#### Cross-validated comparison of computational hypotheses

We tested whether across-trial history improved prediction of behavior beyond current-trial LC stimulation. Three models were compared on the same trials: a controller model containing current-trial WN and the three nuisance covariates defined above; a persistent-state model additionally containing the 20 trial-lag LC-drive history; and a teaching model additionally containing the 20 trial-lag LC-drive x signed-outcome history. Both stimulated and unstimulated current trials were included.

Predictive performance was evaluated using 5-fold outer cross-validation, with identical outcome-balanced folds across models. Within each outer training set, the ridge penalty was selected by fivefold inner cross-validation. The primary measure was Bernoulli log-likelihood on held-out trials; classification accuracy and deviance explained relative to an intercept-only model were calculated as secondary measures. Improvements of the persistent and teaching models were evaluated relative to the controller model on the same held-out folds.

### Temporal coupling between LC activity and behavior

#### LC spiking impulse-response

To account for the temporal transformation between optical stimulation and evoked LC activity, we measured the population spiking impulse response of laser-responsive LC units separately in each monkey. The population impulse response from 0 to 100 ms after optical input was held fixed in the subsequent analyses. We then used a forward model to estimate the timing of LC activity that best accounted for the observed within-trial behavioral kernel. Candidate temporal gates *G*(*t*) in LC activity time were Gaussian functions with centers µ ranging from –200 to 300 ms and widths *σ* from 5 to 200 ms, both in 5 ms increments. Each candidate gate was forward-projected through the measured LC impulse response *h*(*t*) to predict the behavioral kernel as a function of optical-input time,

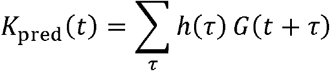

Here, *t* is the time of optical-input relative to sample onset and *τ* is the time after optical stimulation in the measured LC impulse response.

#### Estimation of the LC-time controller gate

Predicted and observed behavioral kernels were compared over the full ™400 to 400 ms interval. Pearson correlation was used to identify the gate center and width that best reproduced the temporal profile of the observed kernel (**Figs. 4j** and **4n**). As a complementary goodness-of-fit analysis, we searched gates containing a constant component,

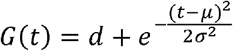

with a signed constant component *d* ranging from –0.4 to 0.4 to capture any sustained offset independently of the transient Gaussian component, and fitted a single behavioral amplitude *a* by least squares, such that

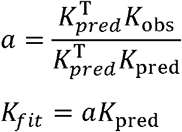

Fit quality was quantified using zero-baseline *R*^2^ as,

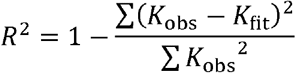

Statistical significance was assessed using 1000 permutations of behavioral outcomes. For each permutation, the behavioral GLM kernel was re-estimated and the complete temporal-gate search repeated, with the maximal fit across the parameter search retained to form the null distribution. Parameter uncertainty was estimated from 1000 bootstrap resamples of behavioral trials, repeating the GLM estimation and gate search for each resample. The measured population LC impulse response was held fixed throughout permutation and bootstrap analyses.

### Single-trial prediction of behavior

Projection-based analyses were performed separately for each monkey and stimulus location to test whether the temporal pattern of LC stimulation predicted behavioral outcome on individual trials (**Fig. 5**).

#### Cross-validated kernel projection

For each stimulated trial, an LC-behavior kernel was estimated from all remaining trials using leave-one-out cross-validation. The kernel was defined as the difference between the mean WN profiles of correct and error trials,

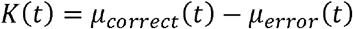

and smoothed with a Gaussian kernel (σ = 15 ms). The held-out trial was projected onto this kernel by taking the dot product between its WN profile and the kernel over an analysis window encompassing the behavioral effect (33% of kernel peak; –110 to –10 ms for monkey S; –140 to –40 ms for monkey P). This yielded one cross-validated projection value per trial, with the held-out trial excluded from construction of its corresponding kernel.

#### Single-trial prediction and ROC analysis

Behavioral discrimination was quantified from the cross-validated projection values using receiver operating characteristic (ROC) analysis and the area under the ROC curve (AUC). Statistical significance was assessed using 5000 random permutations of behavioral outcome labels relative to projection values, with the empirical *P*-value calculated as the proportion of permuted AUC values greater than or equal to the observed AUC. Spearman rank correlation was additionally used to quantify the association between projection value and behavioral outcome, with significance assessed using 10000 random permutations of the outcome labels.

#### Projection-dependent behavioral performance

For visualization of the graded relationship between projection strength and behavioral performance, trials were divided into four equal-sized quantiles according to their cross-validated projection values. The proportion of correct trials was calculated within each quantile.

## Acknowledgments

We thank Dr. Chery J. Cherian and Lai Wei for critical feedback on the manuscript; Dr. Jackson J. Cone and Dr. Mitchell F. Roitman for assistance with optrode fabrication; Morgan L. Bade, Rachel Parker and Autumn O. Mitchell for technical help with monkey procedures.

## Author contributions

S.G. and J.H.R.M. designed the experiments, performed the surgeries, and wrote the paper. S.G. performed the experiments and analyzed the data.

## Data and materials availability

All data are available in the main text or the supplementary materials.

## Supplementary Materials

Supplementary Figures S1 to S7

## REFERENCES

1 Aston-Jones, G. & Cohen, J. D. An integrative theory of locus coeruleus-norepinephrine function: adaptive gain and optimal performance. Annual Review of Neuroscience 28, 403–450 (2005).

2 Poe, G. R. et al. Locus coeruleus: a new look at the blue spot. Nature Reviews Neuroscience 21, 644–659 (2020).

3 Sara, S. J. The locus coeruleus and noradrenergic modulation of cognition. Nature Reviews Neuroscience 10, 211–223 (2009).

4 McBurney-Lin, J., Vargova, G., Garad, M., Zagha, E. & Yang, H. The locus coeruleus mediates behavioral flexibility. Cell Reports 41 (2022).

5 Breton-Provencher, V., Drummond, G. T., Feng, J., Li, Y. & Sur, M. Spatiotemporal dynamics of noradrenaline during learned behaviour. Nature 606, 732–738 (2022).

6 Carter, M. E. et al. Tuning arousal with optogenetic modulation of locus coeruleus neurons. Nature Neuroscience 13, 1526–1533 (2010).

7 Ghosh, S. & Maunsell, J. H. R. Locus coeruleus norepinephrine contributes to visual-spatial attention by selectively enhancing perceptual sensitivity. Neuron 112, 2231–2240. e2235 (2024).

8 Vazey, E. M., Moorman, D. E. & Aston-Jones, G. Phasic locus coeruleus activity regulates cortical encoding of salience information. Proceedings of the National Academy of Sciences 115, E9439–E9448 (2018).

9 AstonLJJones, G. & Cohen, J. D. Adaptive gain and the role of the locus coeruleus–norepinephrine system in optimal performance. Journal of Comparative Neurology 493, 99–110 (2005).

10 Yu, A. J. & Dayan, P. Uncertainty, neuromodulation, and attention. Neuron 46, 681–692 (2005).

11 Grimm, C. et al. Tonic and burst-like locus coeruleus stimulation distinctly shift network activity across the cortical hierarchy. Nature Neuroscience 27, 2167–2177 (2024).

12 Marzo, A., Totah, N. K., Neves, R. M., Logothetis, N. K. & Eschenko, O. Unilateral electrical stimulation of rat locus coeruleus elicits bilateral response of norepinephrine neurons and sustained activation of medial prefrontal cortex. Journal of Neurophysiology 111, 2570–2588 (2014).

13 Todorov, E. & Jordan, M. I. Optimal feedback control as a theory of motor coordination. Nature Neuroscience 5, 1226–1235 (2002).

14 Todorov, E. Optimality principles in sensorimotor control. Nature Neuroscience 7, 907–915 (2004).

15 Bouret, S. & Sara, S. J. Network reset: a simplified overarching theory of locus coeruleus noradrenaline function. Trends in Neurosciences 28, 574–582 (2005).

16 Waterhouse, B. D., Moises, H. C. & Woodward, D. J. Phasic activation of the locus coeruleus enhances responses of primary sensory cortical neurons to peripheral receptive field stimulation. Brain Research 790, 33–44 (1998).

17 Jordan, R. & Keller, G. B. The locus coeruleus broadcasts prediction errors across the cortex to promote sensorimotor plasticity. eLife 12, RP85111 (2023).

18 Glennon, E. et al. Locus coeruleus activity improves cochlear implant performance. Nature 613, 317–323 (2023).

19 Glennon, E. et al. Locus coeruleus activation accelerates perceptual learning. Brain Research 1709, 39–49 (2019).

20 Cone, J. J., Mitchell, A. O., Parker, R. K. & Maunsell, J. H. R. Stimulus-dependent differences in cortical versus subcortical contributions to visual detection in mice. Current Biology 34, 1940–1952. e1945 (2024).

21 Day-Cooney, J., Cone, J. J. & Maunsell, J. H. R. Perceptual weighting of V1 spikes revealed by optogenetic white noise stimulation. The Journal of Neuroscience 42, 3122–3132 (2022).

22 Lehnert, J. et al. Visual attention to features and space in mice using reverse correlation. Current Biology 33, 3690–3701. e3694 (2023).

23 Ghosh, S. & Maunsell, J. H. R. Attention-related modulation in the superior colliculus encodes perceptual sensitivity, but not perceptual choice. Nature Communications 17, 3323 (2026).

24 Ghosh, S. & Maunsell, J. H. R. Single trial neuronal activity dynamics of attentional intensity in monkey visual area V4. Nature communications 12, 2003 (2021).

25 Hautus, M. J., Macmillan, N. A. & Creelman, C. D. Detection theory: A user’s guide. (Routledge, 2021).

