## Supplemental Materials for "Transient noradrenergic signals act as online controllers of perceptual computation"

### Supplementary Figures

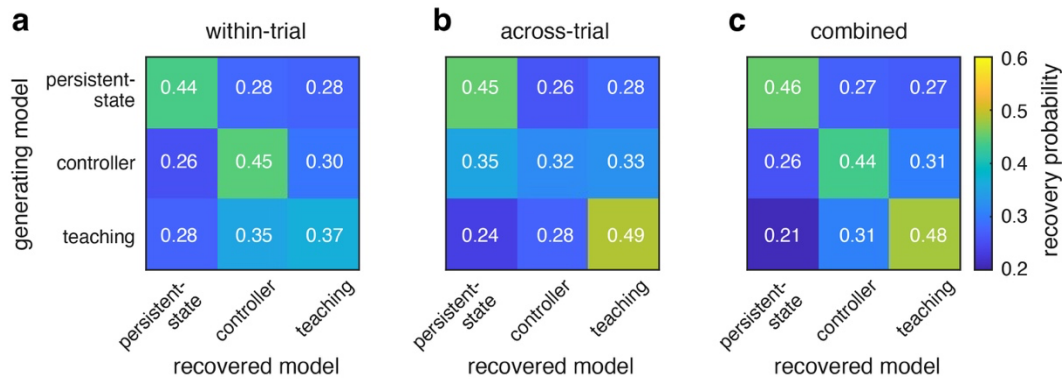

10 **Figure S1. Recovery of the three candidate models from simulated data.** Model-recovery analyses show how the temporal signatures generated by each generating model (rows) could be distinguished from each candidate model (columns) using the computational analysis framework applied in the study. Behavioral choices were simulated separately under each model and each simulated dataset was subsequently fit with all three candidate models. Values indicate recovery probabilities across independent simulations; each row sums to 1, and chance performance is 0.33. (a) Recovery using within-trial perturbation effects. The persistent-state and controller models were recovered with probabilities of 0.44 and 0.45, respectively, whereas the teaching model was less distinguishable from the other models (0.37). (b) Recovery using across-trial effects. The persistent-state and teaching models were recovered with probabilities of 0.45 and 0.49, respectively; the controller model was not reliably identified from across-trial information alone (0.32). (c) Recovery using both within- and across-trial information. Recovery probabilities were 0.46, 0.44 and 0.48 for the persistent-state, controller and teaching models, respectively. Diagonal values indicate correct model recovery and off-diagonal values indicate misclassification. Colors indicate recovery probability. These simulations demonstrate that the complementary within- and across-trial temporal signatures contain information that distinguishes the three proposed mechanisms.

20

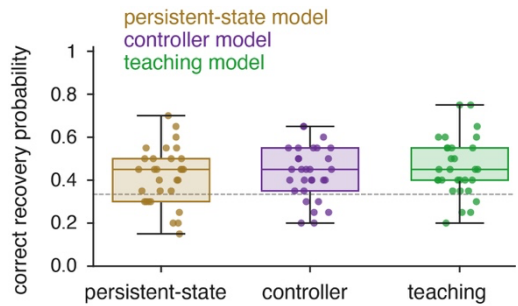

**Figure S2. Model recovery is robust to variation in model parameters.** Recovery of the persistent-state, controller and teaching models was tested across 30 independently sampled parameter sets. For each parameter set, positive gains and time constants were independently sampled between 75% and 125% of their values used in the main simulations. Persistent-state baseline retention and teaching-weight retention were varied additively by $\pm 0.08$

30 and $\pm 0.10$, respectively, around their baseline values and constrained to the intervals 0.55-0.95 and 0.10-0.85, respectively. For the persistent-state model, these included the minimum and maximum state time constants and gains controlling state input, WN input, behavior, sample, memory, test, criterion and lapse effects; baseline state retention was also varied. For the controller model, the control time constant, input gain and sensory gain were varied. For the teaching model, learning rate, behavioral gain and weight retention were varied. Each dot shows the correct recovery probability for one of the 30 parameter sets, calculated from 20 independent behavioral simulations. Box plots summarize recovery across parameter sets (median, 1<sup>st</sup> and 3<sup>rd</sup> quartiles). The dashed line indicates chance recovery (0.33). Recovery was above chance on average for all three models, indicating that model identification did not depend on a single parameter setting.

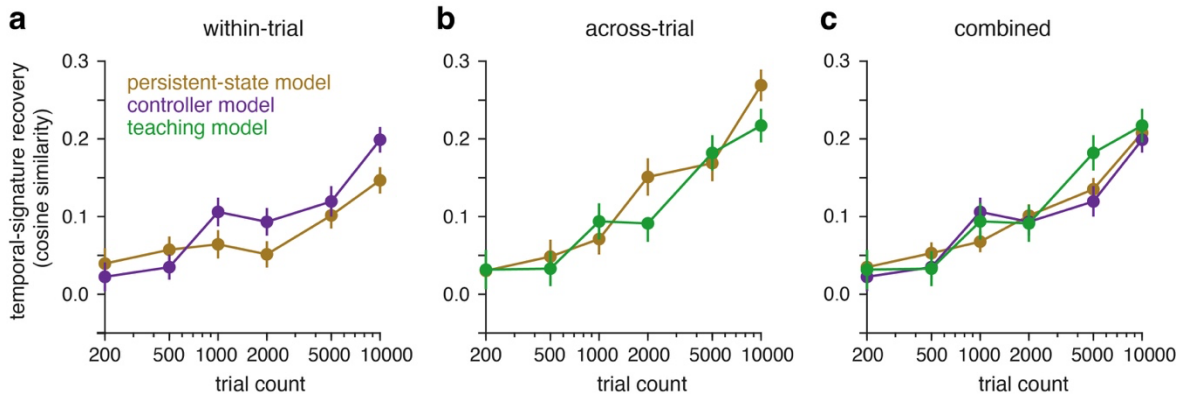

40 **Figure S3. Temporal-signature recovery improves with increasing trial count.** Recovery of the model-specific temporal signatures was evaluated using simulated datasets containing 200, 500, 1,000, 2,000, 5,000 or 10,000 trials. For each trial count, behavioral choices were simulated independently under each model and temporal kernels were estimated using the same analysis applied to the main simulations. Recovery was measured as the cosine similarity between the recovered temporal signature and the corresponding population-limit signature estimated from 50,000 simulated trials. Markers show mean recovery across independent simulations and error bars indicate $\pm 1$ SEM. (a) Within-trial signature recovery for the persistent-state and controller models. (b) Across-trial signature recovery for the persistent-state and teaching models. (c) Combined recovery, summarizing the relevant within- and across-trial signatures for each model. Overall, temporal-signature recovery increased with trial count, showing that larger datasets improved estimation of the characteristic temporal signatures predicted by each model.

50

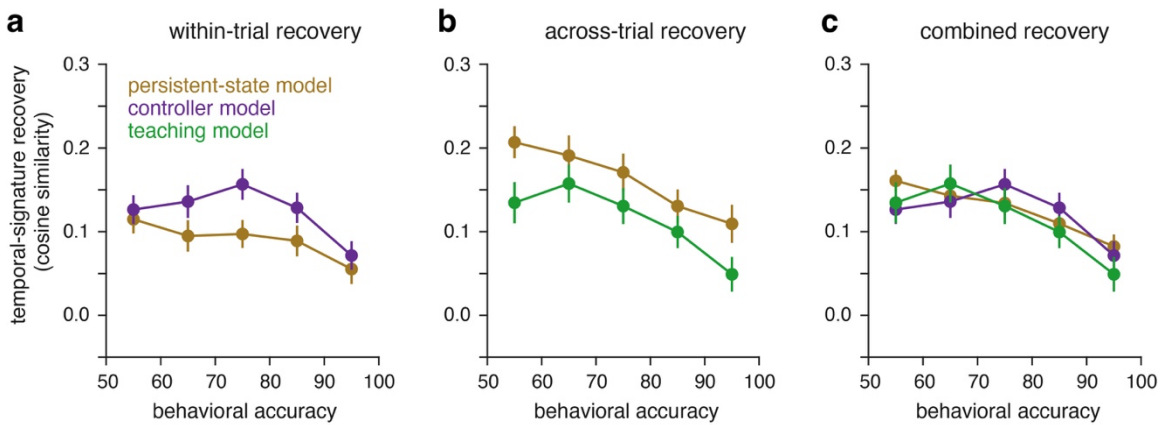

**Figure S4. Temporal-signature recovery across levels of behavioral accuracy.** Recovery of the model-specific temporal signatures was evaluated at behavioral accuracies ranging from approximately 55% to 95%, with the number of simulated trials held fixed at 4,000. Recovery was measured as the cosine similarity between the recovered temporal signature and the corresponding population-limit signature estimated from 50,000 simulated trials. Markers show mean recovery across independent simulations and error bars indicate $\pm 1$ SEM. (a) Within-trial recovery for the persistent-state and controller models. (b) Across-trial recovery for the persistent-state and

60 teaching models. (c) Combined recovery, summarizing the relevant within- and across-trial signatures for each model. Temporal-signature recovery was generally lower at high behavioral accuracy, indicating that recovery depends on behavioral performance even when the number of trials is held constant.

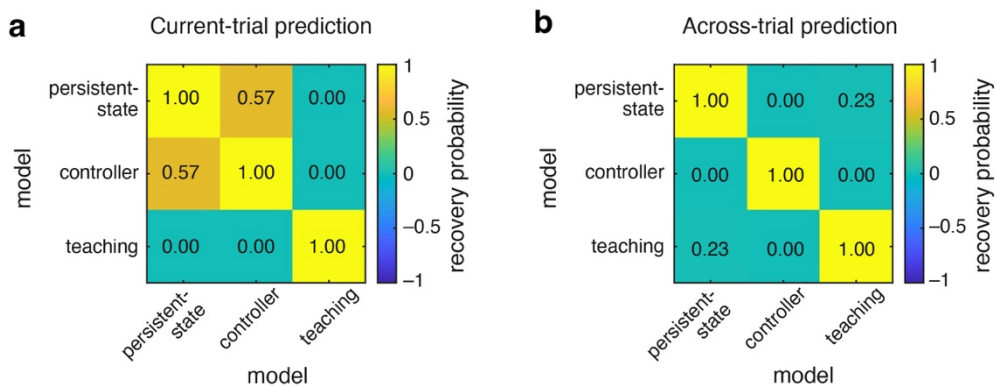

**Figure S5. Similarity of temporal predictions across the three candidate models.** Pairwise cosine similarity was calculated between the temporal perturbation signals predicted by the persistent-state, controller (gate, +100 ms) and teaching models. Larger positive values indicate more similar temporal predictions; diagonal values are 1 by definition. (a) Current-trial predictions. The persistent-state and controller models showed moderate similarity (0.57), whereas the teaching model showed no similarity with either model. (b) Across-trial predictions. The persistent-state and teaching models showed weak similarity (0.23), whereas the controller model showed no similarity with either model. These results show that the three candidate mechanisms produce the expected distinct, but partly overlapping, temporal signatures.

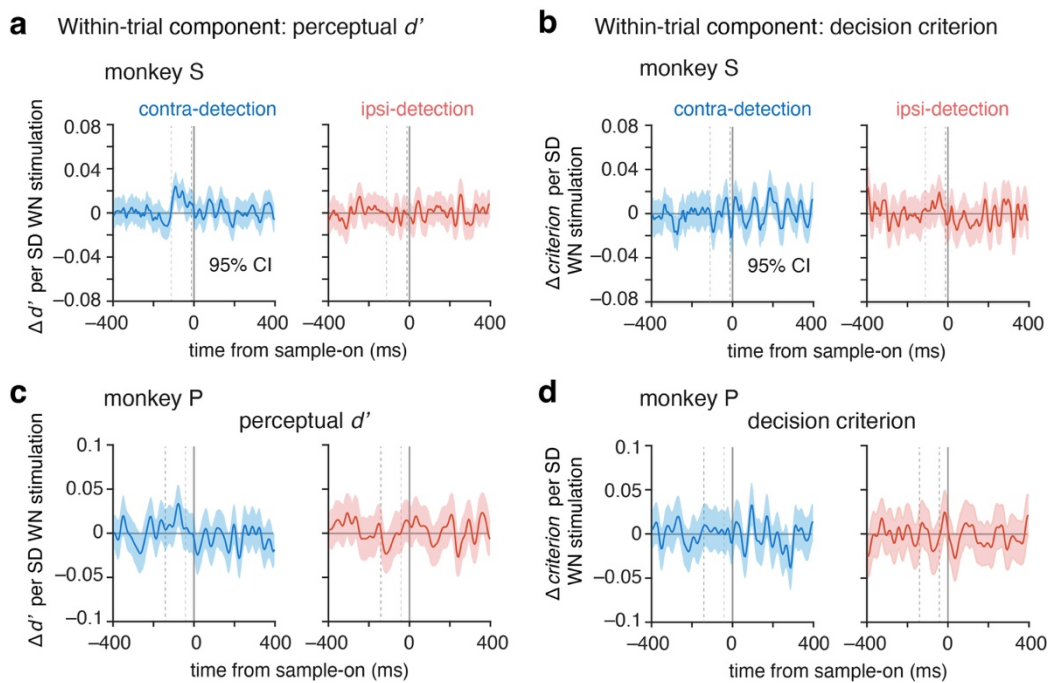

**Figure S6. Temporal kernels of perceptual $d'$ and decision criterion during LC stimulation.** (a, c) Within-trial kernels for perceptual sensitivity ($d'$) in monkey S (a) and monkey P (c). (b, d) Corresponding decision criterion kernels. Kernels were estimated separately for contralateral (blue) and ipsilateral (red) test locations using a ridge-regularized probit model of match/nonmatch reports (Methods). The model included the WN stimulation sequence, its interaction with trial type, and task covariates. Kernel amplitudes express changes in $d'$ or criterion per unit increase in WN input in each 5 ms bin, in standard deviation units. Positive values indicate increased $d'$ or a criterion shift toward reporting 'match', respectively. Shading indicates 95% bootstrap confidence intervals from 1,000 trial resamples. Dashed vertical lines indicate the window used for the window-level SDT analysis.

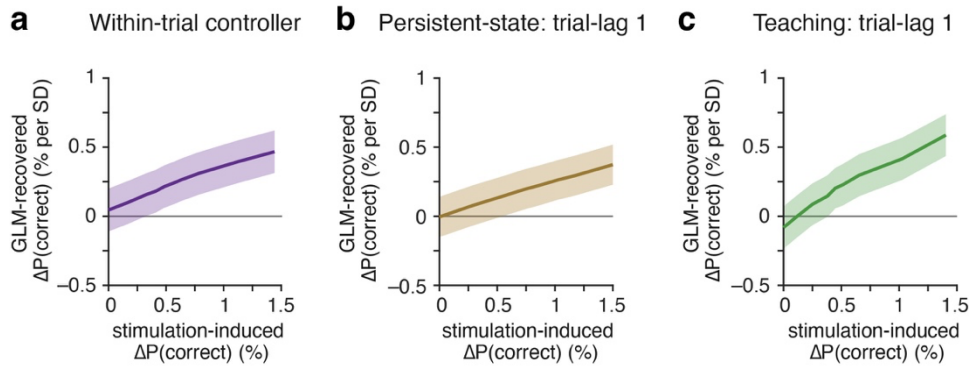

**Figure S7. Sensitivity of within- and across-trial kernel recovery to stimulation-induced behavioral effects.** Sensitivity was evaluated using 200 independent simulated datasets of 2,000 trials each, across a range of generating effect strengths. **(a)** Within-trial controller model, showing recovery of the GLM coefficient for the 0-25 ms white-noise stimulation bin relative to sample onset. **(b-c)** Across-trial recovery of the trial-lag 1 coefficient for the persistent-state (b) and outcome-dependent teaching (c) models. The vertical axis shows the GLM-recovered change in $P(\text{correct})$ per standard deviation of the corresponding predictor. Solid lines indicate mean recovered effects, and shading indicates 95% confidence intervals across independent simulations. The horizontal axis represents the absolute difference in mean $P(\text{correct})$ between matched stimulated and unstimulated conditions, measured on the current trial for the controller model and on the immediately subsequent trial (lag 1) for the persistent-state and teaching models. Kernels were estimated using unregularized logistic GLMs ($\lambda = 0$), with 32 within-trial WN bins or 20 preceding-trial history predictors.

90
